# Complement 3a Receptor mediates high fat diet induced hypothalamic accumulation of lipid associated microglia to regulate neuroinflammation and obesity

**DOI:** 10.64898/2026.04.18.719397

**Authors:** Jean Pierre Pallais, Maria Razzoli, Pedro Rodriguez, Seth McGonigle, Audrey Daugherty, Hannah Hillman, Luca Verteramo, Patricia R. Schrank, Preethy Parthiban, Xuechun Chang, Haiguang Wang, Gianluigi Veglia, Jörg Köhl, Meenakshi Bose, Michelle E. Ehrlich, Stephen R.J. Salton, Alfonso Araque, Daniele Lettieri Barbato, Xavier S. Revelo, Hai-Bin Ruan, Jesse W. Williams, Alessandro Bartolomucci

## Abstract

Microglia, the resident macrophages of the central nervous system, are recognized for their heterogeneity and integral role in brain function and diseases. In the context of high fat diet (HFD) feeding and obesity, microglia become overactive, acquiring a prevailing lipid associated microglial phenotype (also known as LAM). Yet, how microgliosis is induced and regulated remains unclear. Here we report a key role for the Complement 3a Receptor (C3aR), on HFD-induced hypothalamic gliosis and weight gain in mice. HFD consumption leads to elevated microglial expression of C3aR, which parallels widespread accumulation of reactive microglia, selectively in the hypothalamus. Conditional microglial C3aR deletion protects mice from HFD-induced hypothalamic reactive microgliosis. C3aR deletion or pharmacological antagonism opposes HFD-induced weight gain in male but not female mice. Mechanistically, we demonstrated that C3aR is essential for lipid-induced lipid droplet formation, and acquisition of a LAM molecular signature. In summary, we uncovered a previously unknown role for C3aR in the acquisition of a LAM signature driving diet-induced gliosis, identifying this receptor as a new viable therapeutic candidate for conditions associated with hypothalamic neuroinflammation.

## Introduction

Ingestion of high fat diet (HFD) is associated with many inflammation-associated health complications including obesity, type II diabetes, various types of cancers, neurodegenerative disorders, and cardiovascular diseases, which are collectively the leading causes of mortality in the United States and worldwide[1–5]. Hence, there is a need to increase our knowledge on downstream mechanisms informing the development of more effective and targeted therapies[6–9]. HFD feeding and obesity are associated with a state of chronic, low-grade inflammation in many organs, including the brain[10–12]. The hypothalamus, recognized as the homeostatic and autonomic control center, is a primary site of maladaptive neuroinflammation associated with obesity or ingestion of diets rich in fat, ultimately compromising hypothalamic function and leading to metabolic diseases[10–15] and neurodegeneration[16–18] with prolonged exposure.

The brain parenchyma hosts various non-neuronal cell types that are tightly integrated in functional circuits with neuronal populations. The prominent non-neuronal cells in the CNS are astrocytes, microglia, and oligodendrocytes[19,20], with a smaller population of resident immune cells also being present[21]. These non-neural parenchymal cells have gathered growing attention for their still underappreciated roles in health[22–28]. In particular, microglia, the tissue resident macrophages of the CNS, are sensitive to a wide array of nutrients and signals, and capable of transmitting paracrine messengers to nearby neurons and astrocytes to directly regulate brain function and energy homeostasis[24,28,29]. Under HFD and obesity, microglia rapidly expand, particularly in the hypothalamus, which can cause neuronal dysfunction and physiological derangements[10,15,30–32]. If this proinflammatory cascade is persistent, it may become maladaptive, contributing to neurodegeneration, and increasing the risk of developing diseases such as ischemic injury, and Alzheimer’s or Parkinson’s diseases[10,33–36].

While substantial progress has been made in understanding the role of neuroinflammation on disease pathogenesis[14,33,35,36], limited knowledge exists on the molecular drivers of gliosis, besides the established role of fatty acid activation of Toll like receptor 4, and associated proinflammatory response[10–12]. Innate immunity, more specifically the complement system, is likely to play a pivotal role in neuroinflammation and brain diseases due to its early activation in response to various insults[37,38]. Yet, the link between the complement system, gliosis, and neuroinflammation is incompletely understood. Here we report a previously unknown key role for the Complement 3a Receptor (C3aR, encoded by the *C3ar1* gene) on diet-induced hypothalamic gliosis and neuroinflammation. C3aR is a G-protein coupled receptor showing prominent expression in tissue-resident macrophages[39,40] and is the target of two endogenous ligands, the C3-derived anaphylatoxin C3a, and the neural VGF-derived (non-acronymic) peptide TLQP-21[40–45]. The role of C3aR on brain function and neurodegeneration in the context of Alzheimer’s disease is starting to emerge[42,43], but its role in diet-induced neuroinflammation remains unclear. Here we show that C3aR is exclusively expressed by microglia in the CNS, and that consumption of a HFD increases C3aR expression paralleling hypothalamic microgliosis, astrogliosis, and microglial activation, independent of obesity – changes that can be ameliorated by microglial-specific conditional deletion of C3aR. Mechanistically, we found C3aR is a novel marker of lipid associated microglia (LAM) and is required for HFD-induced upregulation of LAM *in vivo* and *in vitro*. Finally, C3aR deletion or pharmacological antagonism opposes HFD-induced weight gain in a sex dependent-manner.

## Methods

### Mouse experiments

All experiments and procedures were conducted with the approval of the University of Minnesota Institutional Animal Care and Use Committee (IACUC, protocol # 2301-40694A). The tdTomato-C3aR1^lox/lox^ reporter knock-in mouse model[39] was obtained in order to localize C3aR expression in the central nervous system. These reporter mice, along with wild-type mice lacking the tdTomato reporter sequence, were used for comparison in the initial characterization of the CNS C3aR. For experiments involving targeted conditional deletion of C3aR in microglia, we first crossed CX3CR1^creERT2/creERT2^ mice (JAX, #:021160) with tdTomato-C3aR1^lox/lox^ mice to generate heterogeneous progeny that were then backcrossed to tdTomato-C3aR1^lox/lox^ mice to generate secondary progeny with necessary experimental genotypes. Following extensive genotyping of the secondary progeny, only those that were homozygous for loxP alleles were used for experiments as they were either or positive for the Cre transgene, resulting in experimental animals with WT (tdTom-C3aR1^lox/lox^;CX3CR1^wt/wt^) or KO (tdTom-C3aR1^lox/lox^;CX3CR1^creERT2/wt^) genotypes once given Tamoxifen (TAM). As this is an inducible system, TAM was dissolved in sterile corn oil (catalog #T5648; Sigma-Aldrich) at a concentration of 20mg/ml and administered via intraperitoneal (IP) injections to these experimental animals (aged 3-4mo) at a dosage of 100mg/kg for five days while housed in their home cages, alternating sides of injection sites each day.

Long term *in vivo* experiments for extensive metabolic characterization involving these C3aR1^lox/lox^-CreER- or C3aR1^lox/lox^-CreER+ animals placed on either a standard diet (Research Diets: SD, D12450B, n=10-12/group/sex) or one of two high fat diets (Research Diets: 45% HFD, D12541 n=10-12/group/sex; 60% HFD, D124192, n=5/group/sex) began nine days after the final TAM injection. Animals were individually housed in static cages and put on a 6AM-6PM light/dark cycle three days to acclimate prior to starting their respective diets. Body weights and food intake were measured twice a week and before starting diets. Body composition was recorded via EchoMRI (EchoMRI LLC, Houston, TX, USA) at baseline prior to starting respective diets and measured biweekly until the end of the experiment. Energy expenditure was measured via indirect calorimetry using the Oxymax Comprehensive Lab Animal Monitoring System (CLAMS; Columbus Instruments) and analyzed using CalR, an online tool for processing metabolic data[46]. Glucose tolerance tests (GTT) were performed following six hours of fasting and measured immediately and up to two hours after intraperitoneal injection of D-glucose (2g/kg). Blood glucose was measured using Accucheck Aviva glucometers (Roche Diagnostics, Indianapolis, IN).

Intranasal delivery experiments (performed completely TAM naïve) were performed as previously described[47], with animals being acclimated prior to commencement of their respective diets and treatment of either sterile saline or DArg10_Aib20[48] (in sterile saline with pH adjusted to 7.4). During the treatment period, animals received 12ul of volume intranasally (3µl bolus in each nostril, two-minute rest, and repeat) daily. DArg10_DAib20 is at a concentration of 8.33ug/ul (3.4mM), for a total of 100µg daily; this concentration is based off of optimal concentrations determined *in vitro*, as well as from pilot dosing experiments performed by our collaborator.

### Cy5-TLQP-21 In Vivo Imaging

Animal care and experimental protocols for in vivo imaging were approved by the Institutional Animal Care and Use Committee at Icahn School of Medicine at Mount Sinai. Seven days prior to imaging, C57BL/6 (Jackson Labs, Bar Harbor, ME, USA) were switched from a standard to an imaging diet [LabDiet 5V75 PicoLab Verified 75/IF (WF Fisher & Sons, Somerville, NJ, USA)] to eliminate interfering autofluorescence. One day prior to imaging, fur on the head and upper abdomen was trimmed, and residual hair was removed by depilatory cream. Fluorescent images were captured using an In Vivo Imaging System (IVIS) Spectrum and LivingImage software (Perkin Elmer, Santa Clara, CA, USA). Mice were anesthetized and placed inside the IVIS imaging chamber with 2% O2 and 2.5% isoflurane delivered via nose cone. Following capture of baseline images, lightly-anesthetized mice were intranasally-administered sterile saline or N-terminal fluorescent Cyanine 5 (Cy5) conjugated TLQP-21 [5 µg/µL; Genscript (Piscataway, NJ, USA)] in saline, with total volumes of 6 µL (30 µg) or 12 µL (60 µg) delivered via pipet gun. Images were captured under anesthesia using excitation λ of 675nm and emission λ of 720 nm (**Supplemental Figure 8**). All images were analyzed for captured radiance (radiant efficiency), and statistical analysis was carried out using Graphpad Prism (Graphpad Software, San Diego, CA, USA).

### qPCRs

RNA was extracted from fresh, microdissected hypothalami using PureLink RNA Mini Kit (ThermoFisher Scientific) according to the manufacturer’s protocol and the concentration was measured using a Nanodrop instrument (ThermoFisher Scientific). Synthesis of cDNA via reverse transcription from isolated RNA was performed using iScript cDNA synthesis kit (Bio-Rad) according to supplied directions. Each cDNA sample was run as a duplicate using iTaq Universal SYBR Green PCR Master Mix (Bio-Rad) to a final volume of 15µL in a CFX Connect thermal cycler and optic monitor (Bio-Rad). Expression of C3aR was normalized to *18S* and *Actin* using BestKeeper[49]. Primer sequences can be found in **Supplemental Table 2**.

### Processing of tissues for *In Situ* Hybridization or Immunofluorescence

Following euthanasia, brains collected primarily for *in situ* hybridization were fresh-frozen on dry ice and stored at -80°C until being embedded in OCT (optimal cutting temperature) compound and then sectioned using a Leica CM1860 cryostat; sections were mounted directly to Superfrost Plus slides (Fisher Scientific) and stored at -80°C until further use. Sections for *in situ* hybridization were sectioned at 10µM and serial sections were collected for immunofluorescence at 30µM, all mounted directly to Superfrost Plus slides and stored at -80°C for later use. For tissues collected only for immunofluorescence, animals were euthanized and exsanguinated by transcardial perfusion with ice-cold PBS for two minutes, followed by perfusion with ice-cold 4% PFA in PBS for 10 minutes as to maintain quality tissue integrity. Whole brain and peripheral tissues were harvested and post-fixed overnight in 4% PFA in PBS at 4°C. The tissues were dehydrated by storage in 30% (wt/vol) sucrose (in phosphate buffer) for a minimum of two days followed by fresh sucrose solution for at least another 24h. Tissues were embedded in OCT and later cryosectioned. Brain were sectioned at 30µM thickness, collected as free-floating sections, and stored in cryoprotectant solution at -20°C. Peripheral tissues were sectioned at 20µM and mounted directly to Superfrost Plus microscope slides (Fisher Scientific) and then stored at -80°C.

### Immunofluorescence

Tissue was washed 3 times for 5 minutes each in PBS (6 washes for free-floating brain sections stored in cryoprotectant). Only brain tissue and not peripheral tissues were permeabilized using 0.3% Triton X-100 in PBS for 15 minutes. Blocking was performed to prevent any nonspecific binding using 5% Normal Donkey Serum (Jackson Immunoresearch; 017-000-121) in PBS for 1 hour at room temperature. Primary antibodies were diluted in blocking buffer and incubated overnight at 4°C, followed by three PBS washes the next morning. Tissue was incubated in blocking solution containing secondary antibodies for 1 hour at room temperature. Tissue was washed 3 times in PBS and then counterstained with DAPI (1µg/ml; Biotium) in PBS for 10 minutes. Sections were briefly rinsed with deionized water and coverslipped using ProLong Diamond antifade mounting medium (ThermoFisher P36970). All imaging was performed on a Keyence BZ-X810 utilizing 10x (N.A. = 0.45), 20x (N.A. = 0.75), or 60x oil-immersion (N.A. = 1.40) objectives. IF imaging at 60x was performed using the optical deconvolution setting to minimize background signal and improve image contrast. Antibody information can be found in **Supplemental Table 3**.

### Immunofluorescence Quantification

For quantification of hypothalamic microglial densities, sections spanning the thickness of the hypothalamus were collected, stained, and sampled [Bregma positions = -0.58mm, -0.94mm, -1.22mm, and -1.70]. 20x stitched images of the hypothalamus were collected at each slice/bregma position, with total Iba1+ cell count (CellCounter plugin) and tissue area manually quantified through FIJI/ImageJ to obtain the cell densities. Iba1+ cell densities were averaged across the four sections per animal. Quantification of the tdTomato fluorescent reporter was performed similarly to that of Iba1+ cells. For astrocyte density, images were converted to grayscale and thresholded to minimize any background or non-specific signal from being quantified. The density of astrocytes was calculated using the measurement function to determine percent area occupied by GFAP+ pixels. Iba1/CD68 and Iba1/Trem2-double positive cells were manually quantified by overlaying the individual image channels in FIJI/ImageJ and using the CellCounter plugin, again with 4 stitched 20x FOVs per animal. Experimenters were blinded to the genotype/sex/diet identifications when processing and quantifying images. Each point represents a sample from an individual animal subject.

### Sholl Analysis for Microglial Morphology

Z-stack images at 1µm step size were collected at 60x magnification utilizing the optical sectioning to achieve highest resolution images necessary for Sholl analysis. The Iba1 stacks were converted to a maximum intensity projection using Keyence BZ-X810 Analyzer and then imported into ImageJ/FIJI for Sholl analysis. Microglial ramification was quantified using a publicly available plugin (neuroanatomy > sholl > legacy: “sholl analysis from image” [set radius = 0-45μm, step = 0.5μm][50]). Three FOVs were collected and processed from each animal (males and females), containing around 5-10 cells/FOV and every cell captured was sampled for analysis. Values from all the microglia per mouse were averaged at each 1µm radial step size from the soma and then AUC was calculated. One-way ANOVA was performed on the AUC values averages across the groups; each point represents the total average of all the individual cells from a given animal subject. Experimenters were blinded to the genotype/sex/diet identifications when processing the images and performing Sholl analysis.

### Fluorescent In Situ Hybridization

RNAscope® fluorescent in situ hybridization (FISH) was performed using RNAscope Multiplex Fluorescent Detection Kit v2 with TSA Vivid Dyes (ACD, catalog#: 323270) according to the manufacturer’s instructions on 10μm coronal cryosections cut from fresh frozen mouse brains. Tissue was first fixed with 4% PFA in PBS for 15 minutes at 4°C and then gradually dehydrated in increasing concentrations of ethanol prepared with milliQ water at room temperature (50%, 70%, and 100% for 5 minutes each). The kit’s “Protease IV” was applied to tissue sections and incubated for 30 minutes at room temperature. Slides were washed 3 times in 1X PBS by placing them in a staining rack and submerging 3-5 times. Slides were then transferred to a HybEZ stain rack (ACD 321716) in a humidified chamber (ACD 310012, 310025), probes were applied and incubated for 2 hours at 40°C in an HybEZ oven (ACD 321710). Probes were all sourced from ACD (**Supplemental Table 4**). Slides were washed 2X2min in the kit’s provided wash buffer in an EZ batch wash tray (ACD 321717). 3 rounds of signal amplification were performed according to manufacturer’s directions – Amp-1 FL for 30 minutes at 40°C, Amp-2 FL for 30 minutes at 40°C, Amp-3 FL for 15 minutes at 40°C. Following signal amplification, Tyramide Signal Amplification reactions were performed by treating slides with FL v2 HRP for 15 minutes at 40°C, followed by incubation with respective TSA Vivid fluorophore for 30 minutes and then FL v2 HRP Blocker for 15 minutes with all steps occurring at 40°C; this process was repeated separately for each individual fluorophore (TSA Vivid 520, 570, or 650 | catalog#: 323271, 323272, 323273). Following the three individual TSA reactions and 2×2min wash, tissue was counterstained with the kit-provided DAPI for 30 seconds. Excess solution was gently tapped off and slides were coverslipped with ProLong Gold Antifade Mountant (Invitrogen P36930). FISH imaging was performed on a Keyence BZ-X810 using the 60x oil objective (N.A. = 1.40) and optical sectioning setting.

### Isolation of primary microglia and immunocytochemistry for tdTom-C3aR1 model validation

Brains of postnatal day P0-3 tdTom-C3aR1^lox/lox^ or tdTom^−/-^ were collected from decapitated pups and the meninges were fully removed. Brains were collected in increments of 3-4 in 6-well plates on ice, each well containing cold Hibernate A media (Catalog number: A1247501; ThermoFisher Scientific). Tissue was homogenized in media using spring scissors, transferred to labeled, sterile 15mL Falcon tubes and then centrifuged at 300g for 5 minutes. The pellets were resuspended in the culture media, filtered through a 70µM nylon cell strainer (Ref: 352350; Falcon) into a clean tube and then seeded in poly-L-lysine pre-coated T75 flasks. Culture media consisted of Dulbecco’s Modified Eagle Medium (DMEM) containing 10% fetal bovine serum (FBS, Invitrogen) and 0.5% penicillin/streptomycin. Mixed cultures were maintained at 37°C and 5% CO_2_. Media was changed the day after dissection to remove dead cells, and then changed every 3 days afterwards. After around 12-14 days, a dense astrocyte monolayer has formed at the bottom of the plate and microglia are loosely attached to said astrocytes. For microglial collection, flasks were agitated at 180rpm for 30 minutes and media was collected and changed for later, repeated collections. Isolated microglia were then plated for immunocytochemistry or calcium influx assays.

For immunocytochemistry, isolated primary microglia (either from tdTom^−/-^ or tdTom^+/+^ animals) were plated to poly-L-lysine pre-coated 8-well slide plates (Part #: 80826, ibidi) and then fixed with 4% PFA in PBS for 15 minutes. Cells were washed with cold PBS and then blocked with 5% Normal Donkey Serum (Jackson ImmunoResearch; 017-000-121) in PBS for 1 hour. Cells were then incubated with a microglial primary antibody (rabbit anti-Iba1, 1:1000, FUJIFILM Wako Chemicals) for 1 hour, washed 3x in PBS, and then incubated with a secondary antibody (donkey anti-rabbit 488; Jackson ImmunoResearch Laboratories) for 1 hour, followed by a final wash series, counterstaining with DAPI, and then imaging.

The calcium influx assay was conducted as previously described ^44,45^. Primary microglia lifted from the astrocyte monolayer were plated to 8-well slide plates and loaded with calcium indicator, 2.5 µM Fluo-4 AM (Invitrogen) for 30 minutes. Cells were washed for 30 minutes with 1x Hanks’ Balanced Salt Solution (Gibco) and then replaced with clean HBSS. The slides were placed on a live cell environmental chamber in a Nikon Ti-E Deconvolution Microscope System with a New Lambda coat anti-reflective coated 20x objective lens. Images of individual wells were captured every second for a total of two minutes. For each recording, the respective individual wells were treated with 10µM TLQP-21 in HBSS just to observe calcium influx. Respective fluorescent images of each well/FOV were captured for tdTom (TxRed) and Fluo4 (GFP). Nikon Elements software was used for the measurement and analysis of the assays.

### Brain Tissue Harvest, Percoll Immune Cell Isolation, and Flow Cytometry

Whole brains were freshly harvested from mice and finely minced using scissors. Tissue was further dissociated by pipetting up and down in 1mL RPMI + 2% FBS, creating the brain suspension. For Percoll gradient separation, 15mL Falcon tubes were layered as follows, from bottom to top: 5mL RPMI + 2% FBS, 1mL brain suspension, 1mL RPMI wash from brain suspension tube, and 3mL 100% Percoll. Warm Percoll to room temperature before use. Tube was gently inverted to mix and 1.5mL 70% Percoll was pipetted using slow drip mode into the bottom of the Falcon tube. Samples were spun with low acceleration and no brakes at 2300rpm for 20 minutes at room temperature. Distinct layers should be visible after spinning. Top myelin layer was vacuumed off, and immune cells at the interface between Percoll and RPMI solutions were pipetted off and transferred into a FACS tube. Isolated immune cells were washed twice in FACS buffer (HBSS + 2% FBS and 2mM EDTA) before proceeding with staining for flow cytometry. Single cell suspensions were passed through a 30μm filter and then spun and resuspended in 50μL of FACS buffer containing antibodies listed in **Supplemental Table 5**. Cells were incubated for 45 minutes at 4°C and then washed twice in FACS buffer. Cells were resuspended in a final volume of 300μL FACS buffer before acquisition on the BD LSRFortessa X-20. Each point represents a sample from an individual animal subject.

### Small Intestinal Immune Cell Preparation and Flow Cytometry

Immune cells were isolated from small intestines following standard protocol, as previously described[51]. For flow cytometric analysis, cells were pre-incubated with anti-CD16/CD32 (Fc block) for 15 minutes on ice to reduce nonspecific binding. For surface marker analysis, approximately 1×10^6^ cells were incubated with fluorophore-conjugated antibodies for 30 minutes at 4°C in the dark, following previously described protocols[52,53]. A fixable viability dye (LIVE/DEAD Fixable Near-IR; Thermo Fisher Scientific) was used at a dilution of 1:500 to exclude dead cells. After staining, cells were washed twice with staining buffer (BioLegend), resuspended in fresh buffer, and maintained on ice until data acquisition. All antibodies/clones used are listed in **Supplemental Table 4**. Flow cytometry data were acquired on a BD LSRFortessa™ (BD Biosciences) equipped with the corresponding lasers and filters for the chosen fluorophores. Acquired data were analyzed using FlowJo software (Tree Star Inc.), where gating strategies were defined using viable single cells and further subdivided based on the expression of relevant surface markers and tdTomato fluorescent reporter. Each point represents a sample from an individual animal subject.

### Primary Microglia Isolation

Following a five consecutive day injection regimen of Tamoxifen (i.p.; 100mg/kg), C3aR1^lox/lox^-CreER- and C3aR1^lox/lox^-CreER+ adult mice were given 9 days to allow for complete depletion before they were sacrificed for primary cell collection. Animals were euthanized via CO_2_ chamber and whole brains were freshly harvested from mice and finely minced using scissors. Tissue was further dissociated by pipetting up and down in 1mL RPMI + 2% FBS, creating the brain suspension. For Percoll gradient separation, 15mL Falcon tubes were layered as follows, from bottom to top: 5mL RPMI + 2% FBS, 1mL brain suspension, 1mL RPMI wash from brain suspension tube, and 3mL 100% Percoll. Warm Percoll to room temperature before use. Tube was gently inverted to mix and 1.5mL 70% Percoll was pipetted using slow drip mode into the bottom of the Falcon tube. Samples were spun with low acceleration and no brakes at 2300rpm for 20 minutes at room temperature. Distinct layers should be visible after spinning. The top myelin layer was vacuumed off, and immune cells at the interface between the Percoll and RPMI solutions were pipetted off and transferred into a FACS tube. Isolated immune cells were washed twice in FACS buffer (HBSS + 2% FBS and 2mM EDTA) before proceeding with magnetic-activated cell sorting (MACS). MACS selection of microglia was performed using the EasySep Mouse CD11b Positive Selection Kit II (StemCell Technologies, Cat.#: 18970) with EasyEights EasySep magnet rack (StemCell Technologies, Cat.#: 18103) in accordance with the manufacturers protocol.

Of these isolated microglia, small aliquots from random samples from each group were (C3aR1^lox/lox^-CreER- and C3aR1^lox/lox^-CreER+) were collected and stained for flow cytometry for validation of successful microglial isolation and terminal confirmation of genotypes. Single cell suspensions were passed through a 30μm filter and then spun and resuspended in 50μL of FACS buffer containing the following antibodies: anti-CD45 BV785 (1:200 dilution, clone: 30-F11, Biolegend #103149), anti-CD11b APC Fire 750 (1:200 dilution, clone: M1/70, Biolegend #101262), and Ghost Dye Violet 510 Viability Dye (1:1000 dilution, Tonbo Biosciences #13-0870-T100). Cells were incubated for 45 minutes at 4°C and then washed twice in FACS buffer. Cells were resuspended in a final volume of 300μL FACS buffer before acquisition on the BD LSR Fortessa X-20.

### Lipid loading & treatments

Cells (BV2 or primary microglia) were cultured in FBS supplemented DMEM, with 100 units/ml of penicillin/streptomycin (Invitrogen, Carlsbad, CA) in a humidified atmosphere of 5%CO2 at 37°C. Once seeded to 12-well plates or Ibidi 8-well slides and allowed to be ∼75% confluent (for BV2 cells), the cells were then treated with 20µg/ml water-soluble cholesterol (Sigma C4951), and 400µM Oleic Acid (Sigma O1257) for 28 hours prior to use for subsequent experiments. Primary cells were plated at a density of 200,000 cells/well.

For the experiments using BV2 cells involving treatment of various antagonists, the cells were pre-incubated with the respective compounds (SB290157 at 20µM, JR14a at 1µM, and DArg10_Aib20 at 20µM) for 1 hour prior to changing to media supplemented with C+O. A separate experiment was repeated with various concentrations of SB290157 or JR14a to establish a dose response. The media that contained C+O also contained the respective antagonists (at the same concentrations) and allowed to incubate for 28 hours. These concentrations were based on previous published work from the lab[44]. The isolated primary cells did not receive treatment with any antagonists, only C+O to induce lipid accumulation and observe if C3aR presence or absence has an effect on lipid droplet handling.

### BODIPY fluorescent imaging and quantification

Following lipid loading as described above, cells plated for imaging were washed, fixed in 4% PFA, counterstained with DAPI and then BODIPY (493/503 at 1:1000 of 1mg/ml working stock; Invitrogen D3922), a stain for neutral lipids. Primary microglial cells were immunostained with a microglial marker for additional validation that those cells were microglia (rabbit anti-Iba1, 1:1000, FUJIFILM Wako Chemicals). All cells were imaged at 60x under a Keyence BZ-X810 fluorescent microscope to visualize the accumulation of lipid droplets as a result of treatment with Cholesterol & Oleate. Fluorescent exposures were kept the same across treatment groups within each cell type. Three FOVs were captured per well.

20x or 60x images of BV2 cells stained with BODIPY were quantified using FIJI/ImageJ. For each FOV, the mean fluorescence intensity (MFI) was calculated using measure function and had the background FI subtracted from that value, and then the corrected MFI was divided by the total number of DAPI nuclei. To obtain the MFI for each well, the values obtained for each of the three FOVs were averaged. An ordinary one-way ANOVA was used to compare the various treatment groups.

For the primary cells, we manually quantified the number of Iba1+ cells and then the Iba1+ cells that were also BODIPY+. Since there were some Iba1+ cells that were not BODIPY+ in both groups, we performed a chi-squared test with Yates continuity correction to show the composition of BODIPY-positive (or negative) in each genotype group.

### Analysis of single-cell RNA sequencing datasets

Data were collected from the human HYPOMAP^50^ and murine HypoMap project[54]. Expression matrices were also collected from publicly available datasets with the following accession numbers: GSE188646[13], GSE208750[55], GSE284492[56]. Raw sequence reads for the PRJNA604055 dataset[57] was downloaded using SRA Toolkit (v3.2.1, https://github.com/ncbi/sra-tools) and aligned using CellRanger[58], with default parameters, using the reference transcriptome of *Mus musculus* (GRCm39).

Cell filtering was conducted according to the following parameters: number of UMIs (>500, <30,000), number of detected features (>400) and percentage of mitochondrial reads (<10%). The scRNAseq analysis was performed in R (v4.5.1, https://www.r-project.org/) using Seurat package (v5.3.0[59]). Data were log-normalized and scaled; highly variable features (HVG) detection (2000 features) and principal component analysis (PCA) were performed. To overcome batch effect given the numerous datasets, data integration was performed using Harmony (v1.4.6[60]). On the integrated objects were conducted non-linear dimensionality reduction using *UMAP* and cell clustering. For visualization, the SCpubr (v3.0.0[61]) package was used.

Cell populations were annotated using common literature markers: Astrocytes (*Slc4a4*, *Slc6a11*, *Itih3*), Microglia (*C1qa*, *Aif1*), Endothelial Cells (*Pecam1*, *Car4*), Neurons (*Bex2*, *Tmem130*, *Ahi1*), Oligodendrocytes (*Mal*, *Mag*, *Trf*), Oligodendrocytes Precursors (*Olig1*, *Olig2*, *Neu4*), Ependymal Cells (*Ccdc153*, *Tmem212*), Fibroblasts (*Pdgfra*, *Col1a1*), Pericytes (*Pdgfrb*, *Rgs5*). The FoamDEX score was determined using the FoamSpotteR module of the MACAnalyzeR package[62], The LAM Score was calculated using the PathAnalyzeR module using a canonical LAM gene signature[63].

### Statistical Analyses

Mice were randomized to diet regimen and for being processed for the various analyses. Investigators were blinded to the genetic identity of the animals from the tamoxifen administration all the way to tissue sample processing and analysis. Data distribution was verified before applying parametric tests. For relative gene expression comparison of *C3aR1*, body weight, percent fat mass content, and hypothalamic density of microglia and tdTom-C3aR1, unpaired t-tests were used. For single time point assessments of %tdTom+ microglia via flow cytometry, unpaired t-tests were used; multiple time points were analyzed with one-way analysis of variance (ANOVA) followed by Tukey’s multiple comparison test. Body weight, body composition, food intake, energy expenditure, GTT, and microglial/astrocyte density were measured and analyzed with two-level, repeated-measures ANOVA followed by Tukey’s multiple comparison test within each sex across diets. If no differences were observed across sexes following 3-way ANOVA, then the data sets were pooled. For glial morphology, total area under the curves were calculated and a one-way ANOVA was performed. For GCaMP6 calcium imaging, unpaired t-tests were utilized across treatments.

## Results

### Selective expression of C3aR by microglia in the central nervous system

C3aR expression is high in the brain, and previous work suggested a prominent expression in microglia[41,42,64,65], but a detailed characterization is lacking. To clarify the cell type specific expression of C3aR in the brain, we took advantage of a knock-in mouse model in which sequence encoding tdTomato (tdTom) fluorescent reporter was fused in frame to sequence encoding the N-terminus of the C3aR protein, with flanking loxP sites for Cre-mediated excision (tdTom-C3aR1^lox/lox^)[39]. After initial observation of diffuse tdTom fluorescence in the brain, including in the hypothalamus (**Fig.S1A-B**) and qPCR confirmation that the fusion of tdTom and insertion of the loxP-flanked cassette does not grossly affect receptor expression (**Fig.S1C**), we utilized cell type markers for microglia, neurons, and astrocytes (anti-Iba1, anti-NeuN, and anti-GFAP respectively) to assign the tdTomato signal to the different cell types. Microglia were the only major cell type in the CNS expressing tdTom-C3aR1 (**Fig.S1D**). Further validation was obtained using RNAscope® fluorescence *in situ* hybridization (FISH). Utilizing probes specific for microglia (*Tmem119*), *C3ar1*, and *tdTom* mRNA confirms the specific expression of *C3ar1* in microglia that colocalizes with *tdTom* (**Fig.S1E**); wild type, tdTom-negative animals had zero *tdTom* signal while *C3ar1* colocalizes exclusively with *Tmem119* (**Fig.S1E**). For tertiary validation of microglial expression of C3aR, primary microglia were harvested from the brains of postnatal day P0/3 tdTom-C3aR1^lox/lox^ and wild type pups. After culturing and isolation of microglia, cells were immuno-stained using anti-Iba1. Fluorescent imaging following immunocytochemistry confirmed that all isolated microglia were tdTom/Iba1 double positive and that primary microglia harvested from wild type mice showed no tdTom fluorescence (**Fig.S1F**). We also confirmed that primary microglia, but not astrocytes that occasionally infiltrated the microglia in culture (which were tdTomato-negative; **Fig.S1G**) from tdTom-C3aR1^lox/lox^ mice respond with a calcium transient in response to the C3aR agonist TLQP-21, in cells loaded with a calcium indicator (Fluo4-AM) (**Fig.S1H**).

Analysis of single-cell RNA sequencing datasets[13,54,56] confirms that microglia are the predominant cell type in the CNS expressing *C3ar1* in mice (**Fig.S2A-C**). *C3ar1* expression overlaps with classical microglia genes, such as *Aif1* and *C1qa* (**Fig.S2C**). This pattern of exclusive microglial *C3ar1* expression was also replicated in a scRNAseq dataset performed on isolated human hypothalami[66] (**Fig.S2D-F**). Unbiased subclustering revealed distinct hypothalamic microglial subtypes in mice (**Fig.1A**) having differing expression of *C3ar1*, along with other LAM-associated genes, such as *Plin2* and *Trem2* (**Fig.1B**). While *Trem2* appears to be broadly expressed by all the 13 hypothalamic microglial subclusters identified, *Plin2* and other LAM markers such as *Ctsb* (Cathepsin B), *Ctsd* (Cathepsin D), and *Cd63*/*Cd68*, are prominently expressed in subclusters 4 and 7; *C3ar1* is also prominently expressed in the same subclusters (**Fig.1B**). MACAnalyzeR analysis tool[62] identified microglial subclusters containing features of foamy macrophages (FoamDEX>0.50). These subclusters were more likely to express *C3ar1*, *Plin2*, *Trem2*, and other LAM markers (**Fig.1C,D**). Similarly, the recently proposed LAM score[63], yields an outcome overlapping to the FoamDEX scores (**Fig.1E**). Ridgeline plots confirm that subclusters 4, 7, 10 align more with that of foamy macrophages/LAMs having high LAM scores (**Fig.1F**). Clusters 4, 7, and 10 are the ones in which *C3ar1* is also expressed the highest (**Fig.1G**). Overall, our analysis identified selective expression of *C3ar1*/C3aR (gene and protein) by microglia, characterizing this receptor as a novel LAM marker in microglia.

**Figure 1.**
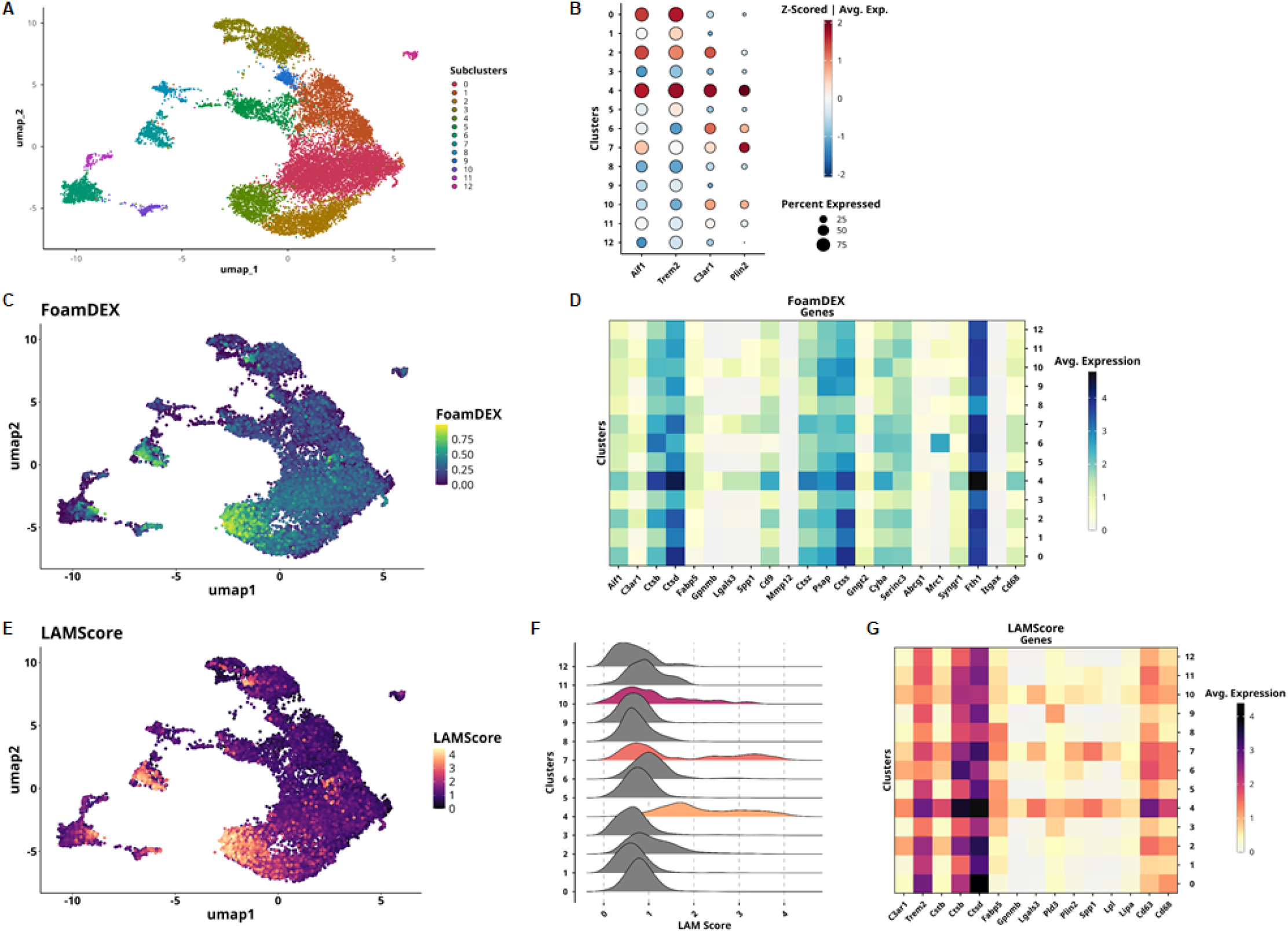
scRNAseq reveals hypothalamic microglial subclusters which are more LAM-like, of which includes C3aR, under physiological, non-obesity conditions. **A)** *UMAP* projection of murine hypothalamus combined scRNAseq datasets showing identified microglial cell subclusters. **B)** Average expression of *Aif1* (Iba1), *C3aR1*,and other LAM genes of interest across the 13 microglial subclusters. C) *UMAP* projection showing FoamDEX score (for identification of foamy macrophages) as determined using MACanalyzeR scRNAseq analysis tools across the various microglial subclusters (FoamDEX score > 0.5 is considered a foamy macrophage). **D)** Heatmap showing average expression of 20 genes across the hypothalamic microglial subclusters that contribute towards the FoamDEX score, with *C3aR1* and *Aif1* included. E) *UMAP* projection showing LAM score, a similar but separate index that mirrors the FoamDEX scores across the subclusters. **F)** Ridgeline plot showing the LAM score across the 13 hypothalamic microglial subclusters, with subclusters 4, 7, and 10 being the most LAM-like. **G)** Heatmap showing average expression of 14 genes across the hypothalamic microglial subclusters that contribute towards the LAM score, with *C3aR1* included.

### HFD feeding causes obesity, hypothalamic microgliosis, and increased C3aR expression

TdTom-C3aR1^lox/lox^ mice were fed either a SD or HFD (60% calories from fat) for twelve weeks. HFD caused elevated body weight gain, percent fat mass, and increased microglial density (microgliosis) of the whole hypothalamus (**Fig.2A-D**), but not of other brain areas such as hippocampus and somatosensory cortex (**Fig.2E-F**). HFD induced microgliosis throughout the entire thickness of the hypothalamus, not localizing to any particular hypothalamic subnuclei (**Fig.2D**). Based on this result, for studies described below, hypothalamic microgliosis was quantified in an anatomically defined position within the hypothalamic PVN, as opposed to quantifying gliosis throughout the full thickness of the hypothalamus. HFD also induced an increase in tdTomato-C3aR expression by microglia (**Fig.2G-H**).

**Figure 2.**
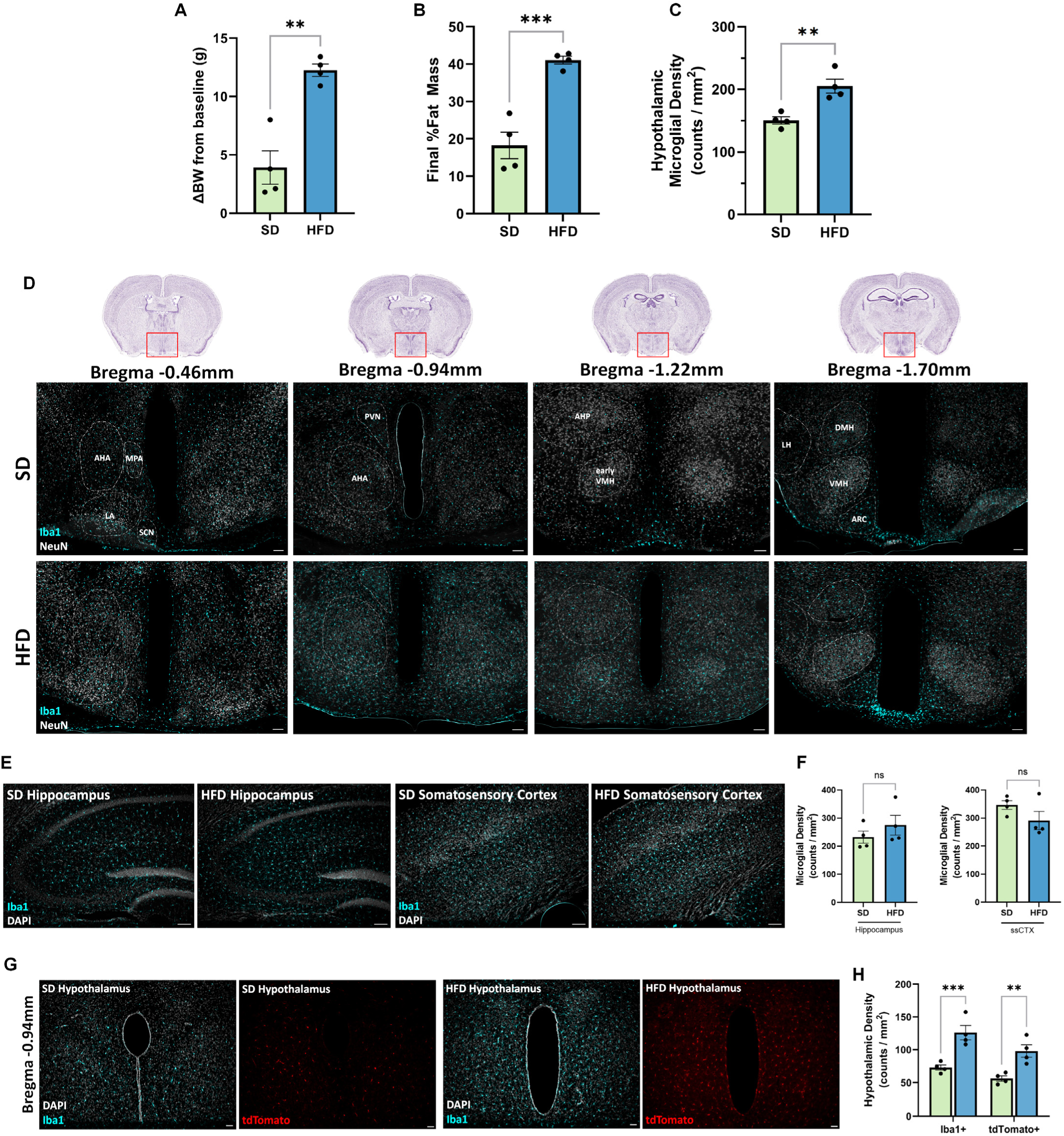
Characterization of diet-induced microgliosis and tdTom-C3aR expression under conditions of diet-induced obesity. 4-month-old male tdTom-C3aR1^lox/lox^ animals placed either on 60% HFD for 12 weeks showing increased body weight gain (**A**; p=0.0016) and fat mass contribution to overall body weight (**B**; %FM/BW; p=0.0008) due to HFD and compared to animals on SD. Unpaired two-sided t-tests; N=4/group. Quantification (**C**) and representative 20x stitched IF images (**D**) showing increased global microglial density (total Iba1+ counts across 4-6 coronal sections/animal divide by total tissue area) spanning the full thickness of the hypothalamus as a consequence of 12 weeks of HFD consumption (p=0.0051). Unpaired two-sided t-tests. Scale bars represent 100µm. Representative 10x IF images (**E**) and quantification (**F**) showing a lack of a HFD effect on microglial density in the hippocampus and somatosensory cortex. Scale bars represent 100µm. Representative 10x IF images (**G**) and quantification (**H**) showing diet-induced hypothalamic microglial density (Bregma -0.94mm, p=0.0039) paralleled by increased tdTom-C3aR expression (p=0.0071). Scale bars represent 30µm. Unpaired two-sided t-tests. ** indicates p<0.01, *** indicates p<0.001. Error bars represent standard error of the mean (SEM).

### Conditional C3aR deletion prevents high fat diet induced hypothalamic gliosis

To test the hypothesis that C3aR is a key regulator of HFD-induced hypothalamic gliosis, we generated a conditional knockout model of C3aR targeting microglia by crossing tdTom-C3aR1^lox/lox^ mice with CX3CR1^creERT2^ mice (Jackson Labs, Stock no.: 021160) to produce tdTom-C3aR1^lox/lox^;CX3CR1^creERT2/wt^ (C3aR1^lox/lox^-CreER+) and tdTom-C3aR1^lox/lox^;CX3CR1^wt/wt^ (C3aR1^lox/lox^-CreER-) and administering Tamoxifen (TAM; **Fig.3A**) at a dosage of 100mg/kg. Mice were fed either SD or HFD diets with increasing level of fat content (45% or 60% calories from fat) for up to 12 weeks. TAM administration didn’t lead to any health complications or significant drops in body weight across any of the groups (**Supplementary Table 1**).

**Figure 3.**
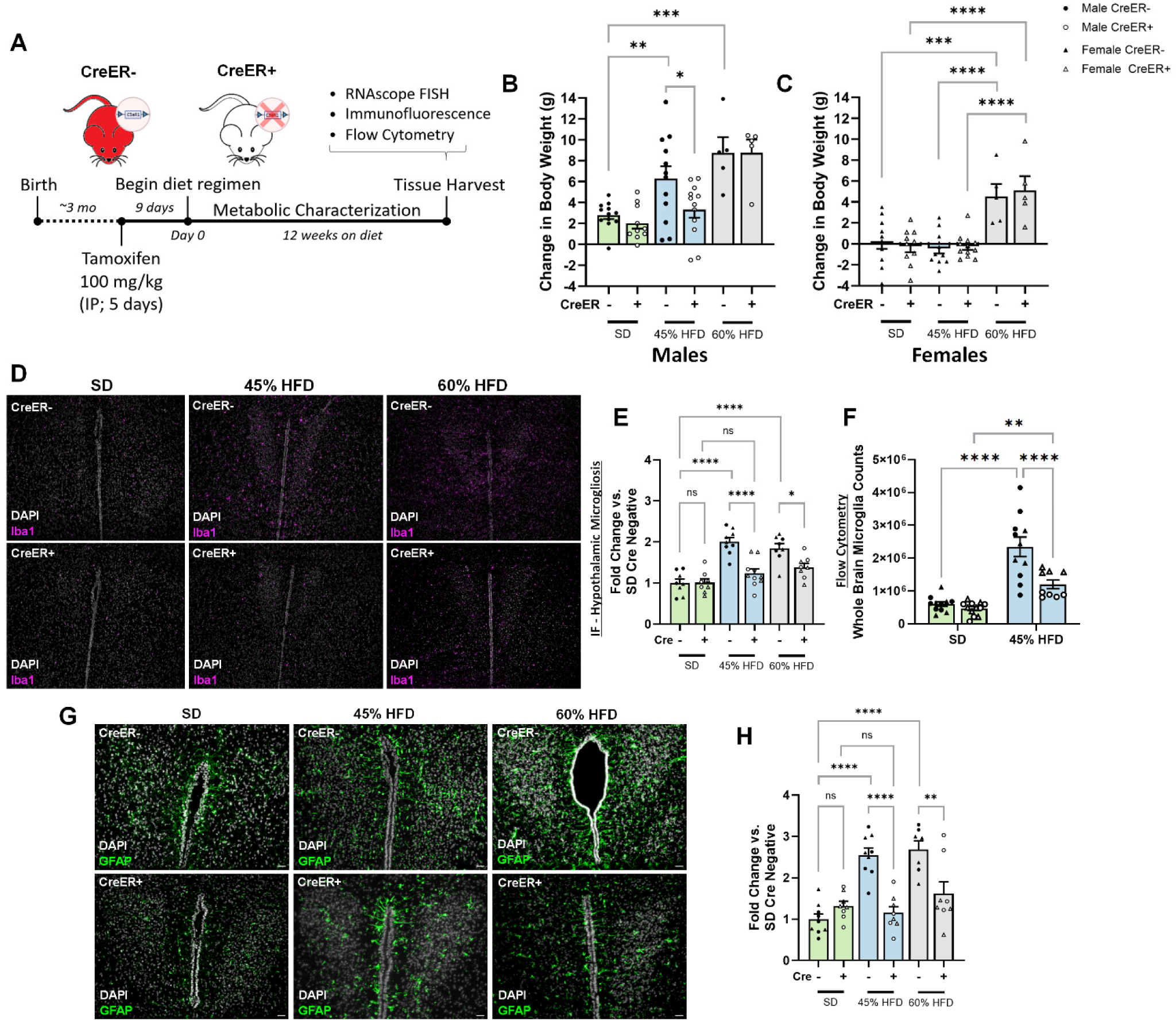
Conditional depletion of microglial C3aR protects against diet-induced obesity and hypothalamic microgliosis. **A)** Experimental timeline for *in vivo* studies. Change in body weight from baseline comparisons within male (**B**) or female (**C**) C3aR1^lox/lox^-CreER- or C3aR1^lox/lox^-CreER+ mice fed either SD, 45% HFD, or 60% HFD for 12 weeks. Male Cre+ animals on 45% HFD were protected against diet-induced weight gain meanwhile female mice regardless of genotype were resistant to 45% HFD-induced weight gain. Males 2-way ANOVA. Diet (main) effect: F(2,50)=18.58, p<0.0001; Genotype effect: F(1,49)=2.415, p=0.1265; Interaction: F(2,50)=1.422, p=0.2507). Females 2-way ANOVA. Diet (main) effect: F(2,47)=26.65, p<0.0001; Genotype effect: F(1,47)=0.02555, p=0.8737; Interaction: F(2,47)=0.3026, p=0.7403). Tukey’s post hoc]. Animals: SD & 45%HFD, N=10-12/genotype/sex; 60% N=5/genotype/sex. Representative 10x IF images (**D**) and quantification (**E**) showing HFD-induced hypothalamic microgliosis that was reduced in animals lacking microglial C3aR. Diet-induced hypothalamic microgliosis was protected when microglial C3aR was ablated in both male and female animals, and regardless of which HFD diet they were on. 2-way ANOVA. Diet (main) effect: F(2,45)=23.12, p<0.0001; Genotype effect: F(1,45)=23.41, p<0.0001; Interaction: F(2,45)=7.515, p=0.0015). **F)** Flow cytometry showing elevated whole brain microglial counts after 12 weeks of 45% HFD that was mitigated in animals with ablated microglial C3aR, consistent with the histology performed on replicate animals in **C/D**. Sexes were pooled although there appears to be clustering based on sex in HFD-fed CreER+ animals (**Fig.S4C**). Pooled 2-way ANOVA. Diet (main) effect: F(1,40)=52.62, p<0.0001; Genotype effect: F(1,40)=9.353, p=0.0040; Interaction: F(1,40)=5.290, p=0.0267). Representative IF images (**G**) and quantification (**H**) showing HFD-induced hypothalamic astrogliosis that was also reduced in animals lacking microglial C3aR, despite these astrocytes not expressing C3aR. Quantification shows an increase in percent area occupied by GFAP+ pixels in 10x magnification images (20x magnification images shown for representative images, **G**). Scale bars represent 30µm. 2-way ANOVA. Diet (main) effect: F(2,43)=15.79, p<0.0001; Genotype effect: F(1,43)=23.34, p<0.0001; Interaction: F(2,42)=13.16, p<0.0001). Error bars represent standard error of the mean (SEM).

Firstly, comparing the change in body weight and fat mass in C3aR1^lox/lox^-CreER-, revealed diet-induced increases in males but not females fed 45% HFD, while both sexes were equally vulnerable to the 60% HFD, concomitant with a significant increase in food intake (**Fig.3B-C; Fig.S3A-E**). 45% HFD caused a significant increase in fat mass gain but a decrease in fat-free mass gain (**Fig.S3E,I**), while 60% HFD caused an increase in both fat and fat-free mass gain (**Fig.S3E,I**). Glucose intolerance measured after 11 weeks on diet was observed only in animals fed 60% HFD (**Fig.S3G-H, K-L**), and no change in energy expenditure was observed when measured after 10 weeks on the diet (**Fig.S4A-D**). Confirming and extending our own previous findings, and data from the literature[10,11,14,34,36], HFD-feeding led to an increase in hypothalamic microgliosis (**Fig.3D-F**) and astrogliosis (**Fig.3G-H**). Both forms of gliosis were independent from sex and percent-fat content in the diet, suggesting that the excess in circulating nutrients and downstream signaling, but not body weight gain itself which was divergent as a function of sex and fat content in the diet, are driving hypothalamic gliosis. Based on these findings, we set out to test the hypothesis that microglial C3aR is causally linked to diet-induced hypothalamic microgliosis and neuroinflammation.

To test this hypothesis, we first validated the experimental mice for conditional C3aR deletion in microglia. Thus, all Cre-mediated deletion of C3aR in the experiments described here is driven solely by the CX3CR1 promoter. CX3CR1^creERT2^ mice are commonly used to target microglia[67–70]. While CreER-activation in postnatal days 3-5 causes microglial activation and aberrant synaptic pruning in this model[71], TAM treatment in adulthood does not result in this adverse effect, while also sparing several peripheral macrophage populations that downregulate CX3CR1 postnatally[69]. Nevertheless, because CX3CR1 is expressed by a subset of macrophages in other organs such as small intestine, liver, and adipose tissue[72], we extended our analysis to other organs as well to confirm specificity of C3aR deletion in microglia long-term. Using IF, FISH, and flow cytometry, we confirmed that TAM administration led to a successful microglial deletion of C3aR in C3aR1^lox/lox^-CreER+ but not C3aR1^lox/lox^-CreER-mice that is detectable as early as two weeks and persists up to 15 weeks post-TAM (**Fig.S5B-C**,**S6E**); C3aR1^lox/lox^-CreER- and TAM-naïve male and female animals retained expression of the receptor (**Fig.S5B**). Firstly, no tdTom was detected in the bone marrow using flow cytometry (**Fig.S6A**), confirming that bone marrow (BM)-derived monocytes are C3aR-negative[39]. Flow cytometry performed on whole brain immune cell isolate, gating for microglia (CD45^Int^; CD11b^hi^) showed that nearly all microglia from TAM-administered C3aR1^lox/lox^-CreER-animals expressed tdTom-C3aR whereas C3aR1^lox/lox^-CreER+ counterparts did not (**Fig.S6B**). Additionally, immunofluorescence and flow cytometry showed that in our experimental condition, tdTom-C3aR1 expression was unchanged in multiple organs analyzed at 2 weeks post-TAM, with the exception of the small intestine (**Fig.S6C-E**). Follow up IHC and flow cytometry experiments confirmed only a transient C3aR depletion in the intestine 2 weeks post-TAM that is fully replenished at later time points (**Fig.S6D-F**). This initial transient depletion of C3aR in peripheral macrophages in the lamina propria of the small intestine may have limited metabolic effect since the expression was normalized afterwards without a noticeable impact on body weight gain. Metabolically, depletion of microglial C3aR protected male mice against diet-induced obesity (only in 45% HFD, not 60%, likely due to exaggerated effects of 60% HFD compared to 45%; **Fig.3B-C**) without any noticeable effect on food intake or glucose tolerance (**Fig.S3B,G,H**). Surprisingly, despite the protection against diet-induced body weight and fat mass gain, there was a slight reduction in cumulative energy expenditure only in male C3aR1^lox/lox^-CreER+ fed 45% HFD (**Fig.S4A-B**), whose functional implication is currently unclear.

Supporting a key role for C3aR in HFD-induced gliosis, deletion of microglial C3aR completely prevented and normalized the diet-induced hypothalamic microgliosis back to SD level in both male and female mice fed either a 45% or a 60% HFD (**Fig.3D-F**). Deletion of microglial C3aR completely prevented the diet-induced astrogliosis as well, normalizing its level back to that observed in SD conditions in both male and female mice (**Fig.3I-J**). This result suggests a generalized beneficial effect of C3aR deletion on glia cells that do not express this complement receptor, such as astrocytes. To extend this analysis, we investigated the impact of C3aR deletion on general neuroinflammation in the brain performing flow cytometry on a separate cohort of male/female C3aR1^lox/lox^-CreER- or C3aR1^lox/lox^-CreER+ animals fed SD or 45% HFD (i.e. the diet in which we observed the more marked effect of the receptor deletion). We first confirmed that there was successful retention/deletion of microglial tdTom-C3aR1 (**Fig.S7A**). We observed a HFD-induced increase in whole brain total cell counts (consisting of immune cells and microglia (**Fig.S7B**)) as well as in microglia (CD45^Intermediate^, CD11b+; **Fig.3F, Fig.S7C**), and non-microglia immune cell counts (CD45^high^; **Fig.S7D**). Conditional microglial C3aR deletion significantly reduced the HFD-induced increase in microglial cell counts in male mice, whereas we observed a trending decrease of total cell counts in females (**Fig.3F, Fig.S7B-C**). This finding is consistent with the reduction in microgliosis as shown via immunofluorescence (**Fig.3D-E)**. Although microglia are the predominant contributing cell type to CNS immune cell isolates, HFD consumption also increased non-microglial immune cell subpopulations such as T cells, B cells, classical monocytes, and neutrophils (**Fig.S7D-H**). Subtle sex differences were observed in how these immune cell subpopulations changed in response to HFD feeding, and C3aR deletion played a minimal role in these changes (**Fig.S7E-H**). Overall, these results suggest that C3aR deletion is crucial for HFD-induced gliosis, while it minimally affects the modest glia-independent neuroinflammation observed in the brain in response to HFD.

### Microglial C3aR deletion prevents high fat diet induced reactive microgliosis

Under pathological conditions such as obesity, neurodegeneration, and other inflammatory or degenerative diseases, microglia not only increase in number but also adopt an amoeboid-like shape, that can be indicative of a proinflammatory state that has been implicated in disease pathogenesis[73]. To investigate the diet-induced reactive inflammatory gliosis, and protection against it due to ablation of microglia C3aR, we took high-magnification images to profile the morphological complexity of microglia[74–76] in mice fed SD or 45% HFD. Sholl analysis revealed that under standard diet and normal C3aR expression (C3aR1^lox/lox^-CreER-fed SD), the hypothalamic microglia had normal ramified morphology, characterized by multiple branches with intricate arborizations (**Fig.4A-C**). C3aR1^lox/lox^-CreER- animals fed HFD exhibited an amoeboid morphology, with simpler branching and reduced arborizations (**Fig.4A-C**). Conditional microglial C3aR deletion in C3aR1^lox/lox^-CreER+ protected against this diet-induced change in microglial morphology in both male and female mice (**Fig.4A-C**). Furthermore, CD68 staining, a proxy of phagocytic activity[73,75,77], further confirm that hypothalamic microglia exist in an activated state in response to HFD, which is reversed upon C3aR deletion (**Fig.4D-E**).

**Figure 4.**
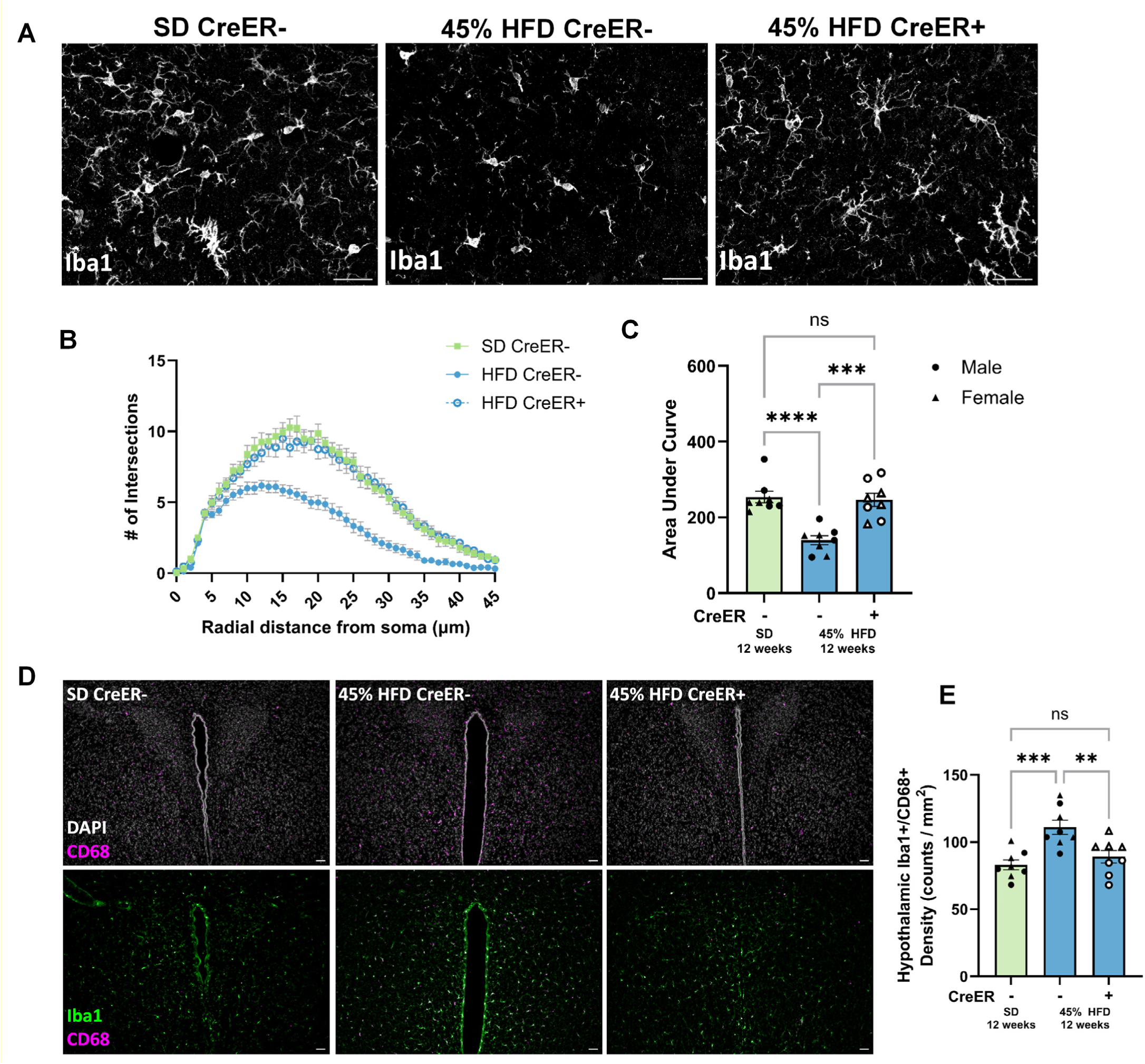
Conditional depletion of microglial C3aR protects against diet-induced changes in microglial activation. **A)** Representative IF images at 60x conveying differences in microglial morphological complexity across diets, notably retracted processes and simplification of arborizations due to 45% HFD that is fully restored in the absence of microglial C3aR. Scale bars represent 30µm. **B)** Sholl analysis displaying the number of intersections at every 1µm radii concentric circle radiating from cell soma, up to 45µm radii circles. **C)** The total area under the curve was calculated per group with male and females pooled due lack of sex differences (sex factor, p=0.1550). Each animal per group is represented by 3 random, averaged fields of view as this is a nested analysis. 1-way ANOVA followed by Tukey multiple comparisons test. *** indicates p<0.001, **** indicates p<0.0001. **D)** Representative 10x IF images showing hypothalamic CD68 expression depending on diet conditions that colocalizes with microglia. There is an observable increase in Iba1 cells colocalizing with CD68 due to diet exposure that is mitigated by deletion of C3aR1, quantification normalized to tissue area shown in **E**. 1-way ANOVA followed by Tukey multiple comparisons test. ** indicates p<0.01, *** indicates p<0.001. Scale bars represent 50µm. Error bars represent standard error of the mean (SEM).

### Intranasal administration of functional C3aR antagonist mimics protective effect of C3aR deletion

To extend the physiological relevance and translational potential of C3aR in HFD-induced obesity and hypothalamic gliosis, we treated 45% HFD–fed male mice intranasally[47,78,79] according to a validated experimental pipeline (**Fig.S8**), with the C3aR functional antagonist, DArg10_Aib20[48], or vehicle for 12 weeks (12µl/d of saline or DArg10_Aib20; 8.3µg/µl), mirroring the conditions in which C3aR conditional deletion conferred a protective effect. DArg10_Aib20 administration limited HFD-induced body weight gain compared to the saline group (**Fig.5A-B**), with a significant decrease in fat mass gain (**Fig.5C**) and no change in fat-free mass (**Fig.5D**), in the absence of changes in food intake (**Fig.5E**). Critically, DArg10_Aib20 administration also protected against HFD-induced hypothalamic microgliosis (**Fig.5F-G**), thus mirroring the effect observed with the genetically driven deletion of the receptor in a pharmacologically relevant approach.

**Figure 5.**
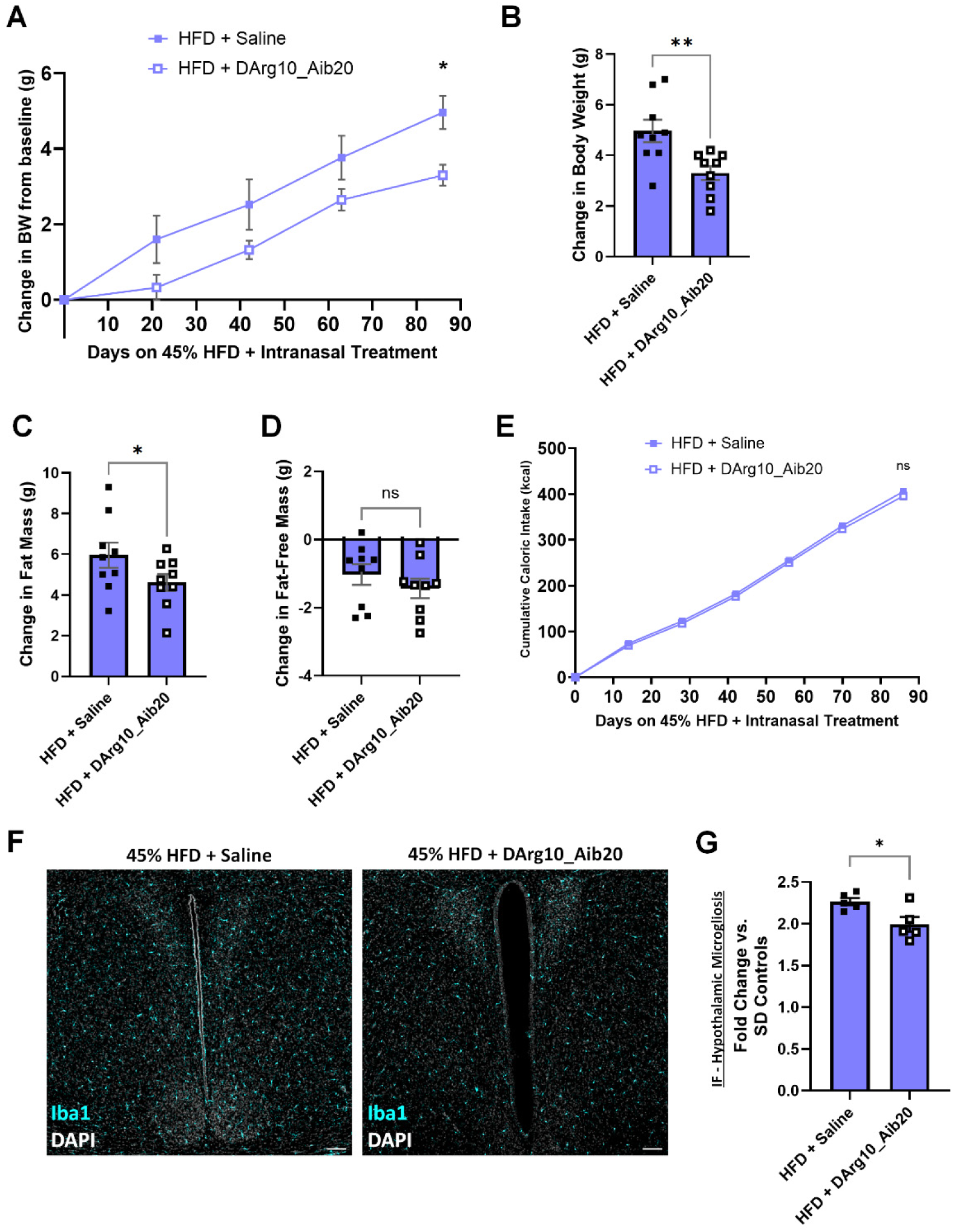
Pharmacological antagonism or deletion of C3aR protects against lipid-induced changes, *in vivo* and *in vitro*. **A)** Change in body weight over time of male mice on 45% HFD receiving daily intranasal administration of saline or DArg10_Aib20 (100µg/day in 12µl). Two-way ANOVA with repeated measures followed by Šídák’s multiple comparisons test (treatment p=0.0392); N=9/group. Final change in body weight (**B**; p=0.0029), fat mass (**C**; p=0.0458), and lean mass (**D**; p=0.1699) relative to baseline of HFD-fed male animals receiving intranasal saline or DArg10_Aib20; Unpaired, one-sided t-tests. **E**) Cumulative caloric intake over time of HFD-fed male mice receiving intranasal saline or DArg10_Aib20. Two-way ANOVA with repeated measures followed by Šídák’s multiple comparisons test (treatment p=0.3235). Representative IF images **(F)** and quantification **(G)** showing HFD-induced hypothalamic microgliosis that was reduced in DArg10_Aib20 treated mice (p=0.0323). 1-way ANOVA (compared to SD fed, control animals) followed by Tukey multiple comparisons test. Scale bars represent 100µm. * indicates p<0.05, ** indicates p<0.01. Error bars represent standard error of the mean (SEM).

### C3aR is a key determinant of lipid-induced LAM

To obtain mechanistic insight into the role of microglial C3aR in conditions relevant to HFD on LAM formation, we first analyzed the expression of Trem2, a marker highly associated with the LAM phenotype[80–82], in the brain of male mice fed 45% HFD, in which a protective effect was observed in the presence of C3aR conditional deletion. As expected, HFD consumption led to an increased hypothalamic density of Trem2+ microglia as determined by immunofluorescence (**Fig.6A-B**). This HFD-induced increase in Trem2 expression was normalized to SD level when microglial C3aR was deleted (**Fig.6A-B)**. Next, we developed a model of lipid loading *in vitro*, using cholesterol and oleic acid. This model was based on previous methods that used one of the two lipids individually[80,83], and based on the observation that cholesterol and oleic acid (C18:1) are the largest lipid components in the HFD and increases substantially compared to SD. BV2 microglia were switched from standard media to media enriched with Cholesterol + Oleate (C+O), for 18 hours. BODIPY staining confirmed significant accumulation of lipid droplets in C+O treated microglial cells compared to untreated microglia (**Fig.S9A**). Consistently, the expression of key LAM genes associated with lipid droplet formation and/or lipid metabolism such as *APOE, Fabp4, HSL, PLIN1, PLIN2,* and LAM marker *Trem2*, were upregulated by C+O treatment (**Fig.S9C**). The expression of *C3ar1* as well as inflammation markers such as *TLR-4, Arg1, CD68*, and *iNOS* were also upregulated by C+O treatment (**Fig.S9D**). Next, we tested the hypothesis that C3aR signaling is required for the LAM phenotype by incubating BV2 with C+O in the presence/absence of one of three ligands showing functional C3aR antagonism, i.e. SB291057, JR14a[84], or our recently developed DArg10_Aib20 [48]. C3aR antagonism prevented C+O-induced accumulation of lipid droplets in BV2 cells (**Fig.6C-D, S9B**), thus supporting a causal role for C3aR signaling in the link between lipid loading and LAM phenotype. To extend this finding, we cultured primary microglia obtained from C3aR1^lox/lox^-CreER- and C3aR1^lox/lox^-CreER+ TAM treated mice (upon confirming C3aR-TdTom deletion via flow cytometry, **Fig.6E**) with media containing C+O for 18h. Staining with Iba1 and counterstaining with DAPI/BODIPY, demonstrate that more than 80% of microglia from C3aR1^lox/lox^-CreER-were lipid loaded. C3aR deletion significantly decreased the percentage of lipid loaded cells to <50% (**Fig.6F-G**). Overall, these results establish a mechanistic requirement of C3aR activity on microglia for lipid droplet formation and a LAM signature.

**Figure 6.**
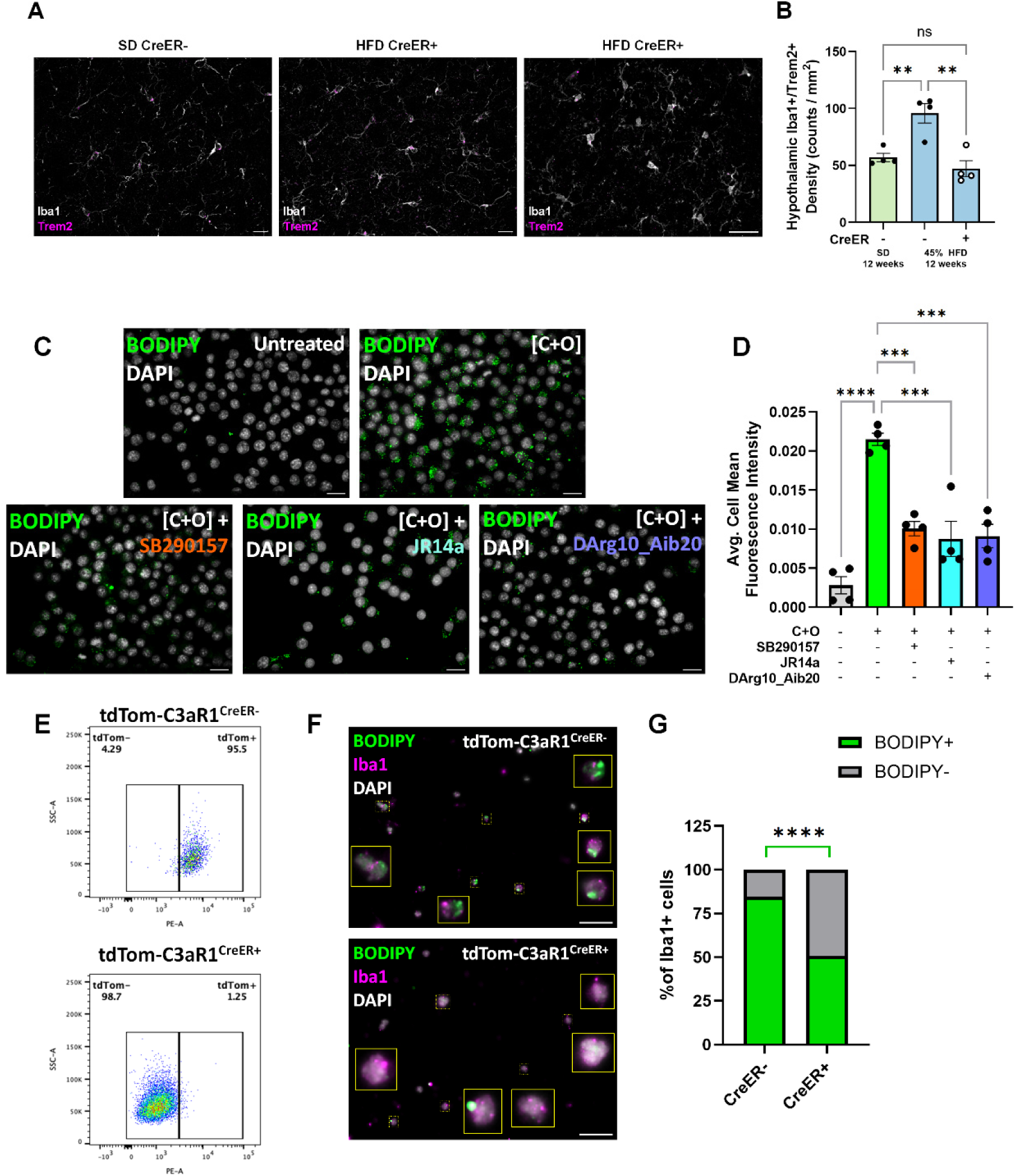
C3aR is important for adoption of LAM phenotype and diet-induced lipid droplet accumulation. Representative images (**A**) and quantification (**B**) showing a diet-induced increase in hypothalamic microglia positive for Trem2 that is protected against in absence of microglial C3aR. 1-way ANOVA followed by Tukey multiple comparisons test. ** indicates p<0.01. **C)** Representative 60x fluorescent images showing lipid droplets in DAPI/BODIPY stained BV2 cells treated with Chol. + OA [C+O] for 18 hours with one of various C3aR antagonists; SB290157, JR14a, or DArg10_Aib20. Scale bars represent 20µm. BODIPY fluorescent intensity quantification normalized to numbers of cells within each field of view for the various groups (**D**). 1-way ANOVA followed by Tukey multiple comparisons test. *** indicates p<0.001, **** indicates p<0.0001. **E)** Representative flow cytometry plot showing identification of tdTom presence/absence in primary microglia isolated from CreER- or CreER+ animals via MACS. **F)** Representative 60x magnification images showing primary cells immunostained with Iba1 and then with DAPI and BODIPY. **G)** Quantification showing a reduction in Iba1+ cells that are also BODIPY+ in primary microglia from CreER+ animals (Chi-square statistic with Yates correction = 16.4987; p-value is 0.000049). Error bars represent standard error of the mean (SEM).

## Discussion

Ingestion of diet rich in fat is an established trigger for hypothalamic neuroinflammation[10,14,85–88], leading to increased risk for significant health complications[36,89], yet the mechanism underlying this process are still unclear. Building on the characterization of its unique expression in microglia, we identified a key role for the C3aR in HFD-induced hypothalamic gliosis, and a sex-specific effect on obesity. Mechanistically, we link dietary lipids to the acquisition of a LAM signature via C3aR. Recent studies implicate C3aR signaling in microglial activation in the context of neurodegeneration, and obesity-related conditions[43,45,48,65]. For example a recent study showed that C3aR signaling adversely affects the ability of microglia to clear amyloid-β plaques, leading to worsened neuropathology and cognitive outcomes[42,43,65,90,91]. Consistently, deletion of C3aR exerts a protective effect that restores normal microglial function and ameliorates associated cognitive deficits in AD mouse models through improved clearance of amyloid-β plaques[43,65,90]. These studies identified a central role for C3aR in microglia function and dysfunction in neurodegeneration, yet its role in diet-induced neuroinflammation remains poorly characterized.

### An essential role for C3aR in HFD-induced hypothalamic gliosis

Diet-induced hypothalamic gliosis can occur rapidly, often within days and can eventually affect other regions of the brain as well over extended periods of time[15,30,92,93]. Upon HFD exposure, we observed a strong increase in the expression of C3aR paralleling hypothalamic microgliosis, which was independent of sex and body weight gain. The notion that hypothalamic inflammation and gliosis are primarily driven by HFD consumption rather than body weight gain is consistent with previous data[14,30,85,94]. Conditional deletion or pharmacological antagonism of microglial C3aR normalized HFD-induced hypothalamic gliosis, suggesting that microglial C3aR is a major contributor in diet-induced expansion of glial cells, but also that the differentiation of BM-derived monocytes (which are C3aR negative) into Iba1+ macrophages is minimal under the present experimental condition, suggesting that local proliferation is the primary source of additional microglia/macrophages in this context[95]. We showed that microglial C3aR not only affects hypothalamic microglial density but also morphology, which is indicative of pro-inflammatory activation[74–76]. In response to HFD and many other pathogenic conditions, microglia acquire a proinflammatory state characterized by a change in shape and branching, and acquiring an amoeboid form[16,22,30]. As expected, microglia from mice fed HFD showed a reduction in their morphological complexity, presenting with amoeboid-like somas and retracted processes and arborizations. Similarly, HFD resulted in increased CD68 staining as a proxy of phagocytic activity[73,75,85]. C3aR deletion reverted the diet-induced amoeboid-like structure and normalized CD68 staining in mice fed HFD irrespective of obesity and sex. The excess of circulating nutrients or lipid-driven metabolic signaling, as well as elevated C3a production from activated astrocytes [96], which together promotes microglial expansion and activation [97–100] may account for the HFD-induced microgliosis.

### C3aR exerts a sex dependent role in HFD-induced obesity

C3aR, and its endogenous ligands has been implicated previously in obesity[101–105]. However, a mechanistic understanding of how C3aR activation by microglia exerts a weight modulating effect is lacking. Here we showed that in the context of HFD consumption, the sex-independent hypothalamic microgliosis was paralleled by significant obesity in a sex-specific manner. When mice were fed a 45% HFD, only males significantly gained weight, while a 60% HFD was necessary for both sexes to become obese. Notably, C3aR deletion protected male but not female mice from 45% HFD-induced obesity. The relative obesity resistance in females was possibly masking a protective metabolic effect of C3aR deletion, and further studies are needed to elucidate the mechanism of this sex-dependent vulnerability, especially since neither food intake measured during the entire study or energy expenditure assessed after 10 weeks on the diet were affected by gene deletion. Nevertheless to extend this finding, we treated 45% HFD-fed male mice with daily intranasal administration of the synthetic C3aR functional antagonist, DArg10_Aib20[48]. Once bound to C3aR, DArg10_Aib20 is biased towards β-arrestin signaling showing otherwise minimal agonist effect, leading to internalization of the receptor limiting agonist mediated activation[48]. DArg10_Aib20 protected against diet-induced weight gain and hypothalamic microgliosis, an effect that recapitulates the beneficial effect of conditional C3aR deletion. This result highlights the potential of developing CNS-restricted selective C3aR antagonists[44,48,106] as a therapeutic strategy to limit hypothalamic inflammation that is associated with a beneficial metabolic effect (at least in male mice), having the potential to exert protective effects on other conditions also associated with brain gliosis such as cognitive impairments, psychiatric and neurodegenerative disease[36,107–109].

### C3aR is a new LAM marker, necessary for lipid-induced lipid droplet formation

Mechanistically, we established for the first time a role for C3aR as a marker, and a potential key regulator of the LAM phenotype. There is a growing appreciation of the role that LAM and lipid droplet accumulation plays in microglia in metabolic and neurological diseases, as well as in aging[110–112]. *APOE*[113,114] and *Trem2*[81,82,110]*, but also PLIN1/2*, *HSL*, and *Fabp4* are among the most common LAM markers[62,63]. Here we established C3aR as a novel marker of LAM using both snRNAseq profiling and functional validation *in vivo* and *in vitro*. Not only can C3aR be considered as a novel LAM marker, but it appears to be a determinant of the increase in expression of LAM genes and lipid droplet formations in response to excess nutritional lipids. Indeed, we observed a HFD-induced increase in the density of microglial cells positive for *Trem2,* suggesting an adverse effect of Trem2 LAM on hypothalamic inflammation associated diseases[81,115,116]. The HFD-induced increase in the density of Trem2 LAM is fully prevented by C3aR ablation, positioning this complement receptor upstream of Trem2 in LAM differentiation. Furthermore, we showed that essential components of the HFD, such as cholesterol and oleic acid, caused lipid droplet accumulation and an inflammatory and LAM-like signature in microglia *in vitro*, including increased C3aR expression. Conditional deletion, or pharmacological inhibition of C3aR by DArg10_Aib20[44], blocked the accumulation of lipid droplets, demonstrating that this receptor plays an essential role in the link between dietary lipids and the LAM signature. Analysis of publicly available single-cell RNA sequencing datasets[13,54,56] revealed that even under healthy, physiological circumstances, there are unique sub-populations of hypothalamic microglia exhibiting a LAM signature. Of all those microglial subclusters, it was these LAM-like microglia that had the highest enrichment of C3aR, alongside other classic LAM markers like *Trem2* and *Cd68*. Further sequencing studies in the context of diet-induced obesity, looking at how these hypothalamic microglial subclusters shift relative to healthy conditions, is needed.

## Conclusion

Given the detrimental impact of diet-induced hypothalamic inflammation on health[10,18,36,107,117–119], understanding the mechanisms underlying this complex phenomenon is essential. Here, we demonstrate that microglial C3aR expression parallels diet-induced hypothalamic gliosis and a LAM phenotype, and that targeted conditional ablation or pharmacological antagonism provides a protective effect. The studies presented here highlight the potential for developing and utilizing selective antagonists of C3aR as a therapeutic strategy to preserve CNS function under conditions associated with hypothalamic gliosis and diet-induced neuroinflammation, reducing the risk and incidence for associated health complications and disease.

## Acknowledgements

This work was supported by NIH/NIDDK R01DK117504 (AB), NIH/NIDDK R01DK102496, (AB), Institute for Diabetes Obesity and Metabolism University of Minnesota (2022 Pilot and Feasibility Program) (AB), Minnesota Partnership for Biotechnology and Medical Genomics (#24.01; AB), IBP Grant Accelerator Program (AB), NIH/NIA R01AG084699 (MEE and SRJS), NIH/NIA R01AG093879 (MEE and SRJS), Cure Alzheimer’s Fund (MEE and SRJS). JPP and PR received support from NIH/NIDDK T32DK083250. HH was supported by American Heart Association pre-doctoral fellowship [25PRE1361476]. JWW was supported by NIH/NIAID R01AI165553 and MN Dept. of Education Award (#214654). We’d like to thank the Research Animal Resources at the University of Minnesota for their instrumental role in animal care on a daily basis. We’d also like to thank the University Imaging Centers and the staff for their training and invaluable support in executing the various calcium imaging experiments. We also thank G. Friedman for her assistance in tissue processing, histology, imaging, and image processing. We also thank Dr. S. Graves and Dr. D. Baker for generously sharing expertise in the Sholl analysis of glial morphology.

## Data availability

All data will be made publicly available upon acceptance of the paper.

## Author Contributions

JPP, MR, PR, SM, AD, HH, PRS, PP, XC, HW, MB performed experiments under the supervision of XSR, S-BR, JW, MEE, SRJS and AB. LV analyzed single-cell RNA sequencing datasets under supervision of DLB. JK and GV provided reagents. JPP and MR performed statistical analyses. JPP and AB drafted the manuscript that was approved by all authors. AB conceived the study and was responsible for funding.

