## Supplementary Figures and Tables for "Complement 3a Receptor mediates high fat diet induced hypothalamic accumulation of lipid associated microglia to regulate neuroinflammation and obesity"

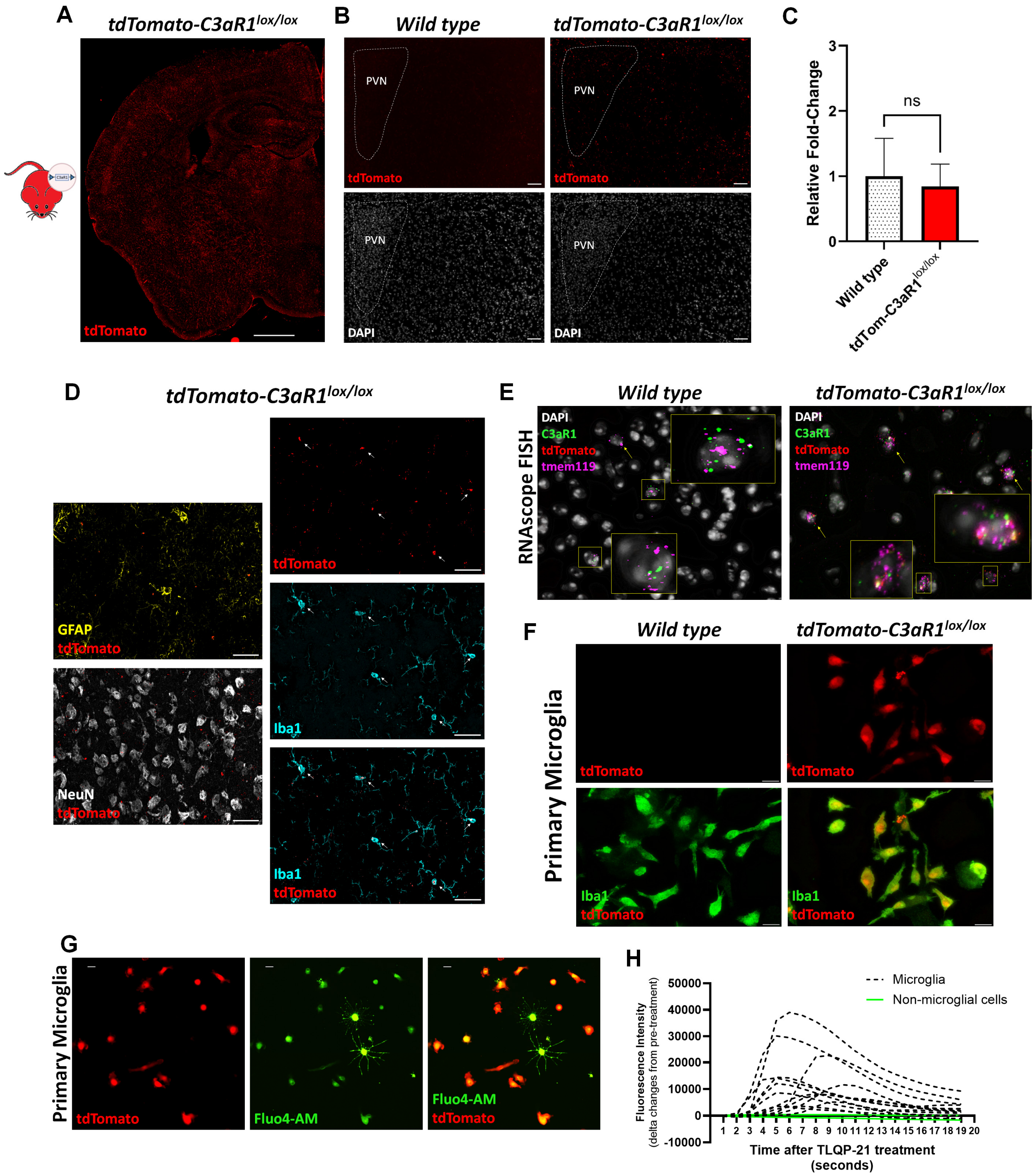


**Supplementary Figure 1. Characterization of C3aR expression in the central nervous system (CNS). A)** Representative 10x stitched image of a half-coronal section from tdTomato-C3aR1$^{lox/lox}$ animal showing broad tdTomato fluorescence throughout parenchyma. Scale bar represents 500µm. **B)** Representative 20x IF images comparing tdTomato autofluorescence between wild type and tdTom-C3aR1$^{lox/lox}$ animals near paraventricular nucleus (PVN) of hypothalamus. Scale bars represents 50 µm. **C)** Relative-fold change in gene expression of C3aR1 in microdissected hypothalamus from wild type or tdTom-C3aR1$^{lox/lox}$ animals, showing no difference in C3aR1 expression due to genotype (N=8/group). Error bars represent standard error of the mean (SEM). **D)** Representative 60x images showing a lack of tdTomato colocalization with neurons (NeuN) or astrocytes (GFAP), only to be observed in microglia (Iba1). Scale bars represent 30µm. **E)** Representative 60x of fluorescence *in situ* hybridization between wild type and tdTom+ animals showing additional confirmation of C3aR1 transcripts being found on microglia (Tmem119), with tdTomato transcripts only being present on tdTom-C3aR1$^{lox/lox}$ mice. **F)** Primary microglia isolated from wild type mice stained for Iba1 exhibited no tdTom autofluorescence, whereas iba1+ primary microglia isolated from tdTom-C3aR1$^{lox/lox}$ animals were tdTom+. **G)** Representative images showing mixed glial cells loaded with calcium indicator (Fluo4-AM) from tdTom-C3aR1$^{lox/lox}$ mice. Astrocytes showed zero tdTom autofluorescence adjacent to tdTom+ microglia. **H)** Fluo4-AM loaded mixed glial cells treated with C3aR1 agonist, TLQP-21 (10μM) only led to increases in fluorescence intensity in microglia (yellow traces) but not astrocytes (blue traces). Error bars represent standard error of mean (SEM).

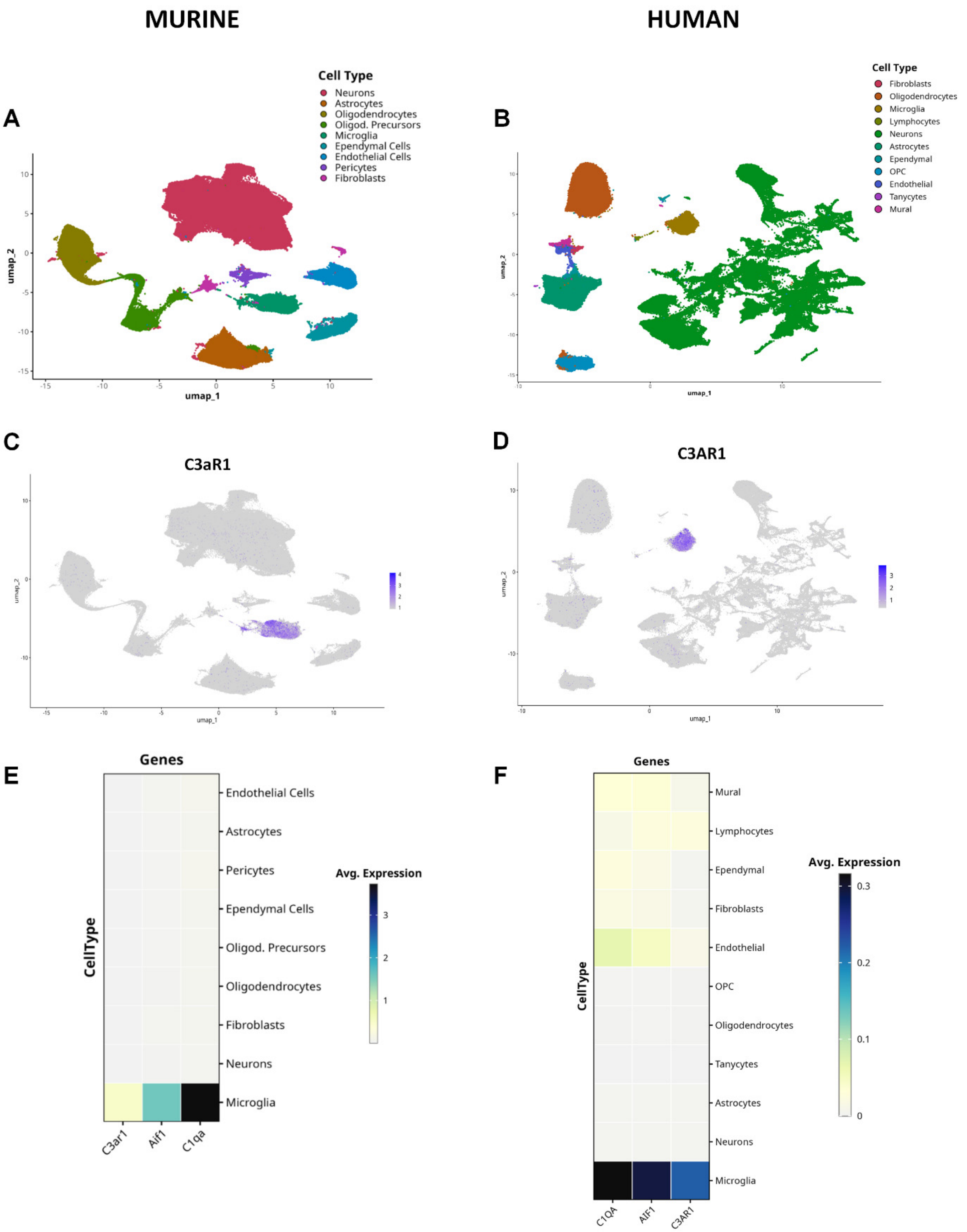


**Supplementary Figure 2. Published scRNAseq datasets validate exclusive, microglial expression of C3aR.** *UMAP* projection of murine (**A**) or human (**B**) hypothalamus combined scRNAseq datasets showing identified cell clusters. *UMAP* projection showing the expression of *C3ar1* gene across clusters in murine (**C**) or human (**D**) hypothalamus. Heatmap of gene expression of microglia markers (*C1qa*, *Aif1*) and *C3ar1* in hypothalamus of mice (**E**) or humans (**F**) grouped by cell populations.

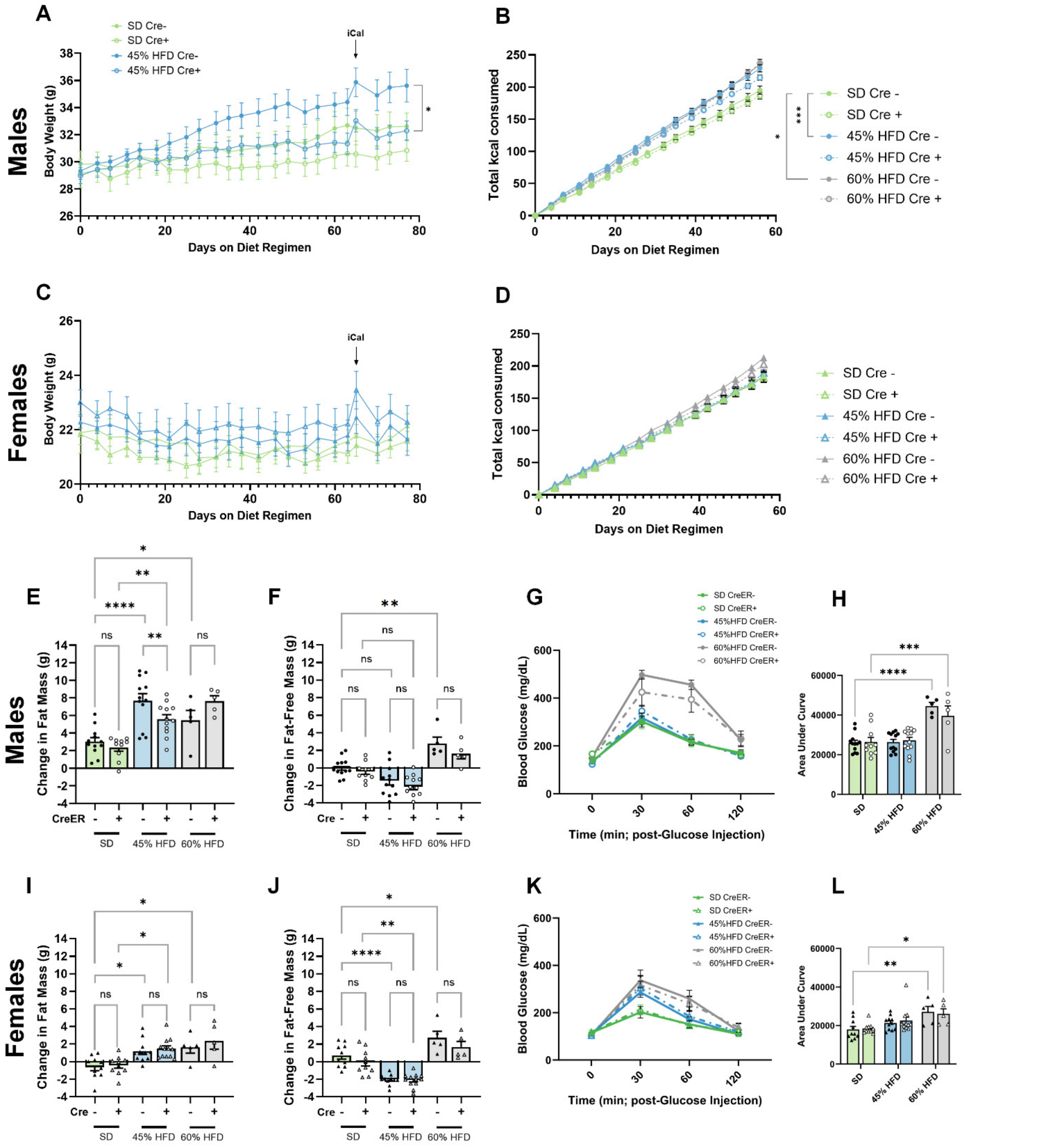


**Supplemental Figure 3. Metabolic characterization after conditional *C3aR1* gene deletion.** Body weight trajectories and cumulative food intake over time for TAM-administered C3aR1$^{lox/lox}$-CreER- or C3aR1$^{lox/lox}$-CreER+ males (**A-B**) and females (**C-D**) fed SD or 45% HFD for 12 weeks. Additional metabolic measures, including changes in body composition and glucose tolerance test for males (**E-H**) and females (**I-L**). Male C3aR1$^{lox/lox}$-CreER+ animals fed HFD showed a significant reduction in diet-induced fat mass accumulation, similar to their overall body weight gain phenotype (**A**). Animals on 45% HFD lost fat-free mass, regardless of sex/genotype (**F,J**). Glucose tolerance test following 6 hours of fasting showing no differences in baseline fasting glucose or glucose handling between male (**G-H**) or female (**K-L**) C3aR1$^{lox/lox}$-CreER- or C3aR1$^{lox/lox}$-CreER+ animals on SD or 45% HFD; only 60% HFD fed animals showed worsened glucose handling also shown by area under the curve calculation. Males 2-way ANOVA. Diet (main) effect: $F(2,50)=25.95$, $p<0.0001$; Genotype effect: $F(1,50)=0.4336$, $p=0.5133$; Interaction: $F(2,50)=0.8248$, $p=0.4442$). Females 2-way ANOVA. Diet (main) effect: $F(2,47)=9.648$, $p=0.0003$; Genotype effect: $F(1,47)=0.03812$, $p=0.8460$; Interaction: $F(2,47)=0.1813$, $p=0.8348$). Error bars represent standard error of the mean (SEM).

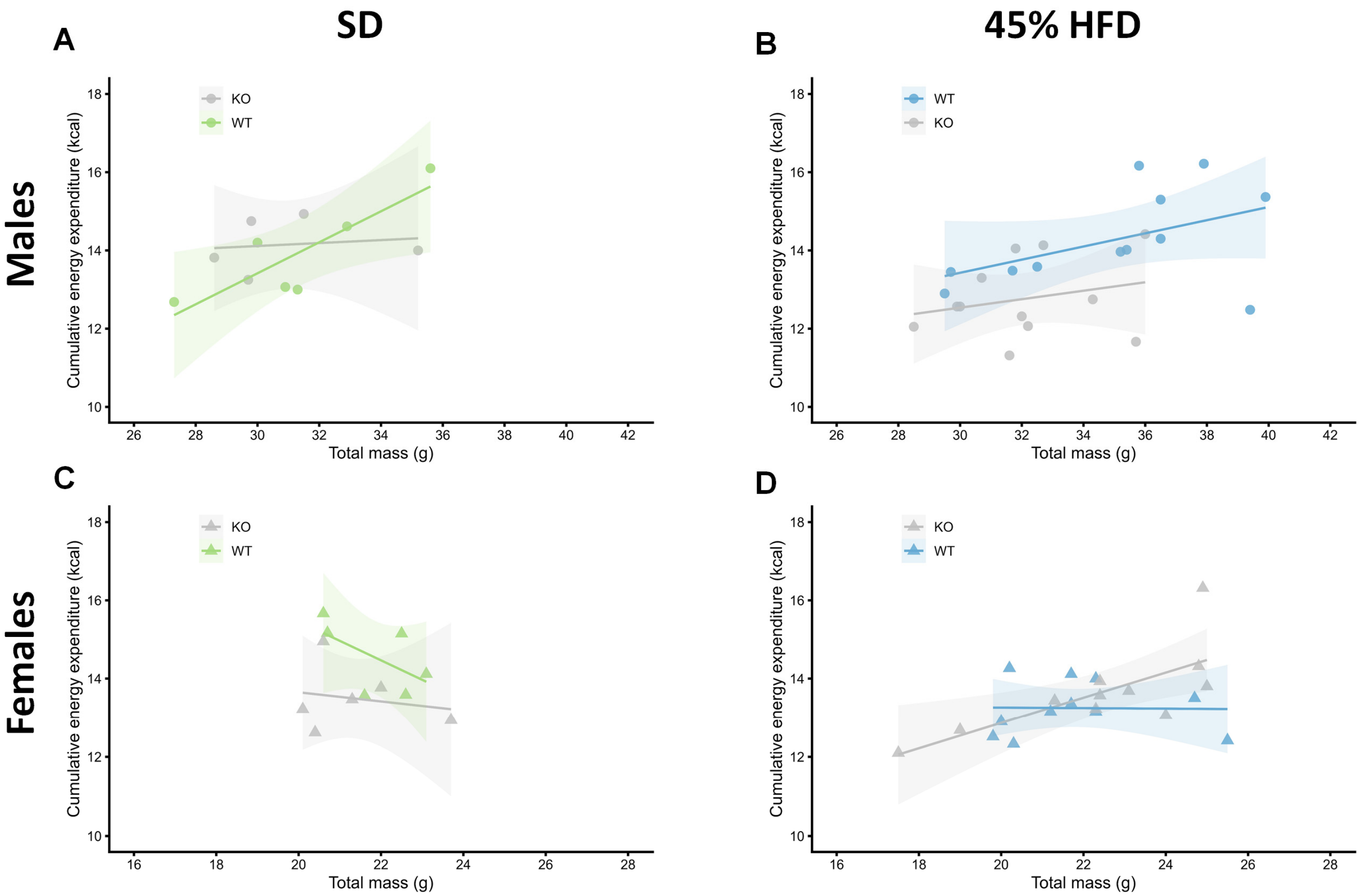


**Supplemental Figure 4. Further metabolic characterization, showing energy expenditure with total body mass as covariate.** Comparisons of cumulative energy expenditure, with energy expenditure (total kcal) as the dependent variable versus total body mass (g) as the independent variable. Regression plots for male C3aR1$^{lox/lox}$-CreER- or C3aR1$^{lox/lox}$-CreER+ on SD (**A**) or 45% HFD (**B**). Regression plots for female C3aR1$^{lox/lox}$-CreER- or C3aR1$^{lox/lox}$-CreER+ on SD (**C**) or 45% HFD (**D**).

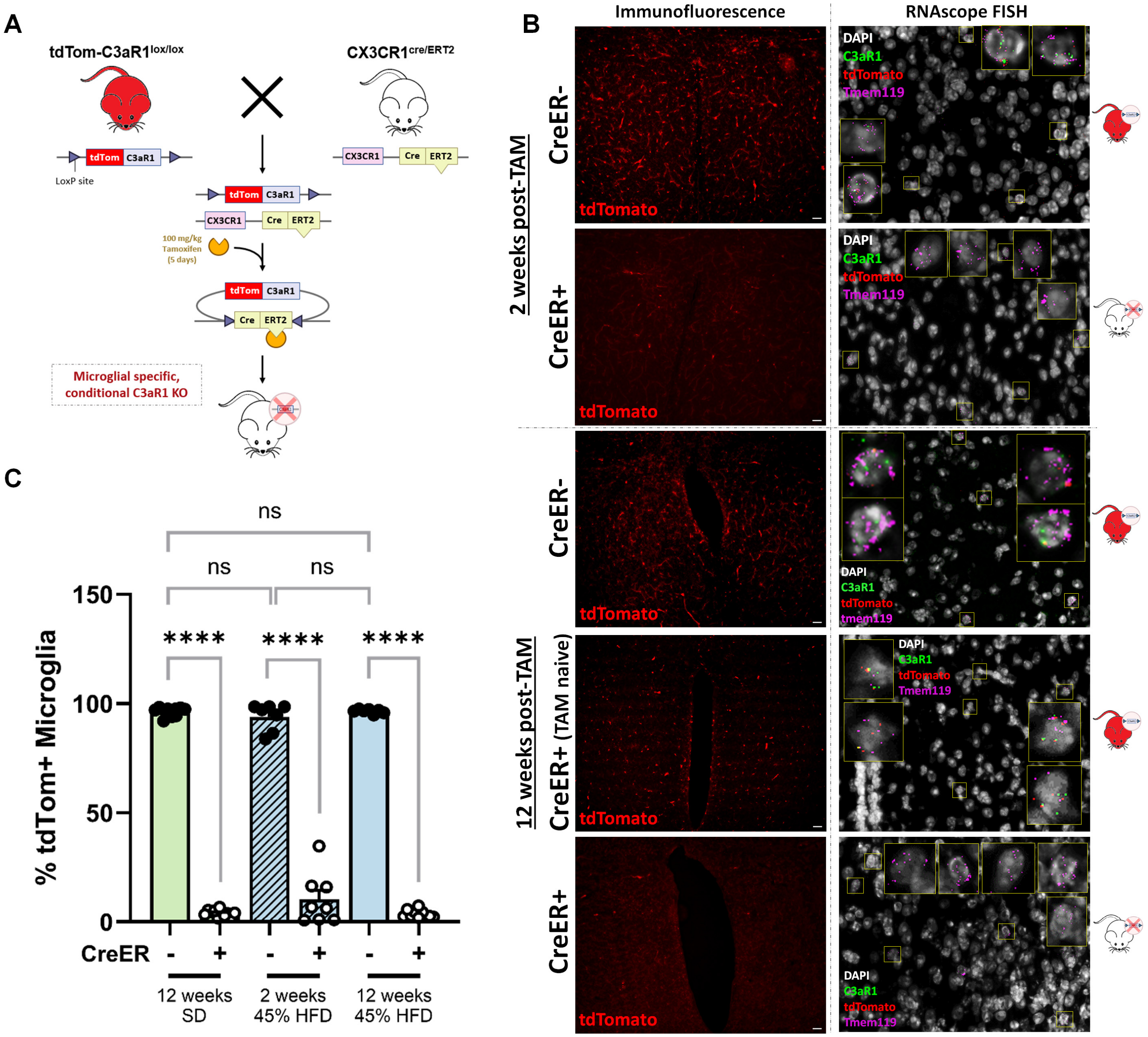


**Supplemental Figure 5. Generation and validation of a persistent inducible, microglial C3aR knockout mouse model. A)** Scheme to generate inducible knockout model targeting microglial C3aR using Cre-recombinase driven by CX3CR1. **B)** Representative images of IF (20x) and FISH (60x) performed on serial sections from TAM-administered C3aR1$^{lox/lox}$-CreER- / C3aR1$^{lox/lox}$-CreER+ mice, or TAM-naïve C3aR1$^{lox/lox}$-CreER+ animals 2 or 12 weeks post-TAM. **C)** Flow cytometry performed on whole brain immune cell preps collected from TAM-administered C3aR1$^{lox/lox}$-CreER- or C3aR1$^{lox/lox}$-CreER+ pooled male/female animals either on SD (12wks post-TAM; N=11-12/group) or 45% HFD [2wks (N=7-8/group) or 12wks (N=7-10/group) post-TAM] showing complete depletion of microglial C3aR that persists up to at least 12 weeks with no loss of deletion efficiency due to HFD. 2-way ANOVA. Genotype (main) effect: $F(2,49)=4.903$, $p<0.0001$; Time effect: $F(2,49)=0.0227$) Tukey's post hoc]. Histogram bars represent group mean. **** indicates $p<0.0001$ Error bars represent standard error of the mean (SEM).

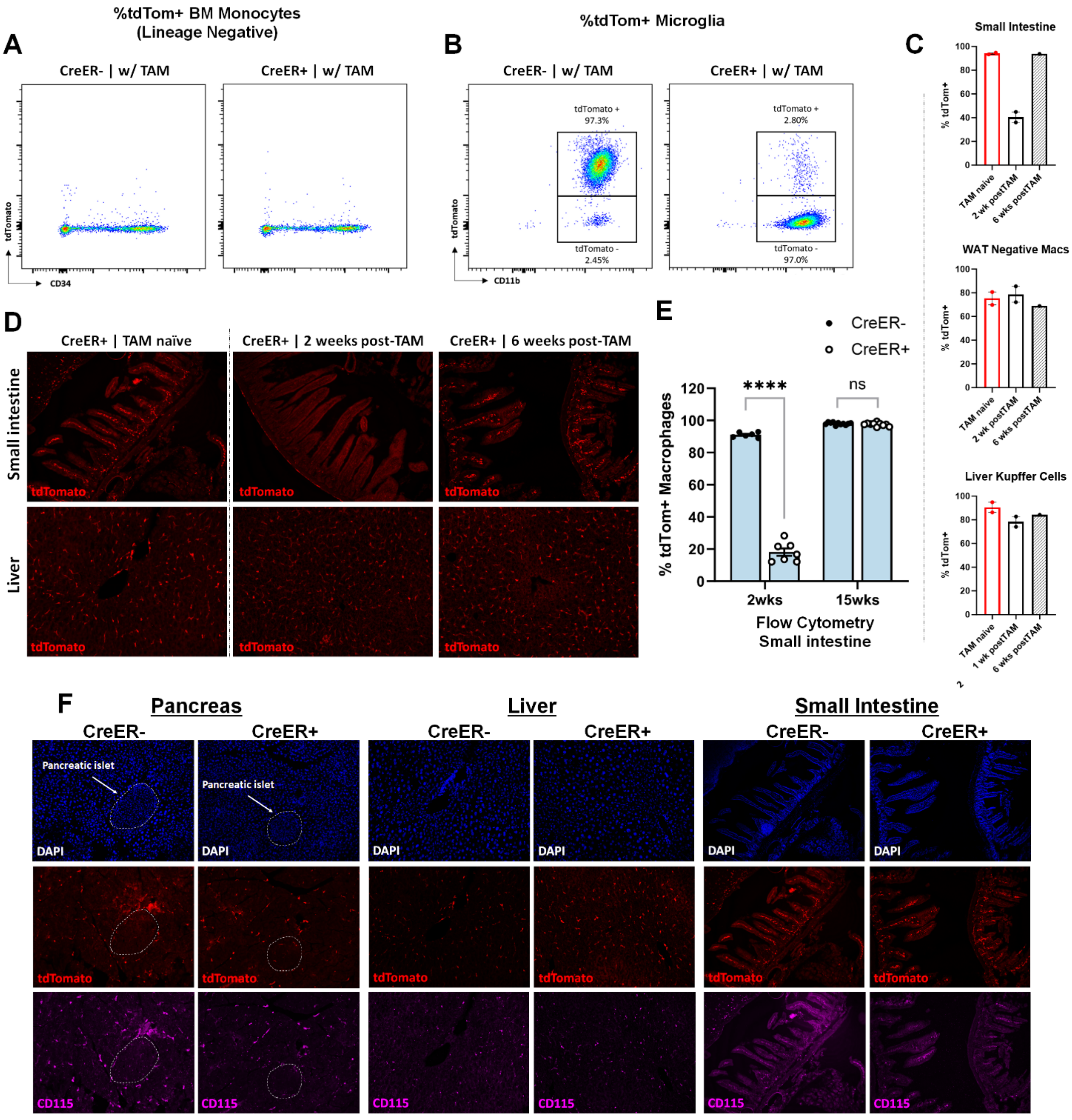


*All peripheral tissues shown were collected 12 weeks post-TAM*

**Supplemental Figure 6. Additional validation and peripheral characterization of C3aR expression after conditional gene deletion. A)** Representative flow cytometry gating of bone marrow-derived monocytes (lineage negative, CD34) showing complete lack of tdTomato expression in both TAM-administered C3aR1$^{lox/lox}$-CreER- and C3aR1$^{lox/lox}$-CreER+ animals. **B)** Representative flow cytometry gating showing high percentage of tdTom+ microglia in TAM-administered C3aR1$^{lox/lox}$-CreER- and near complete abrogating in TAM-administered C3aR1$^{lox/lox}$-CreER+ animals. **C)** Percentage of tdTomato expressing peripheral resident macrophages from small intestine, white adipose tissue, and liver collected from C3aR1$^{lox/lox}$-CreER+ animals 1 or 6 weeks post-TAM, compared to TAM-naïve C3aR1$^{lox/lox}$-CreER+ control. Only peripheral depletion of tdTomato is observed in small intestine, guiding further peripheral characterization. **D)** Representative IF images showing transient depletion of tdTom-C3aR1 only in the small intestine and not liver of TAM-administered C3aR1$^{lox/lox}$-CreER+ animals that is fully reobtained by six weeks after TAM. **E)** Flow cytometry performed on isolated small intestinal immune cell preps from the same cohort of animals shown in **Fig.3C**. Again, transient tdTom-C3aR1 depletion in C3aR1$^{lox/lox}$-CreER+ mice is observed and fully replenished at later time points. **F)** Representative IF images of metabolically relevant pancreas, liver, and small intestine from TAM-administered C3aR1$^{lox/lox}$-CreER- or C3aR1$^{lox/lox}$-CreER+ (animals shown in **Fig.4**) showing lack of any deletional effect in pancreas and liver and no long term effect in small intestine; 12 weeks post TAM. **G**) Indirect calorimetry in TAM-administered in male/female C3aR1$^{lox/lox}$-CreER- and C3aR1$^{lox/lox}$-CreER+ animals 2- or 12-weeks post-TAM to determine possible effects of transient peripheral C3aR depletion on energy expenditure; no differences were observed. Error bars represent standard error of the mean (SEM).

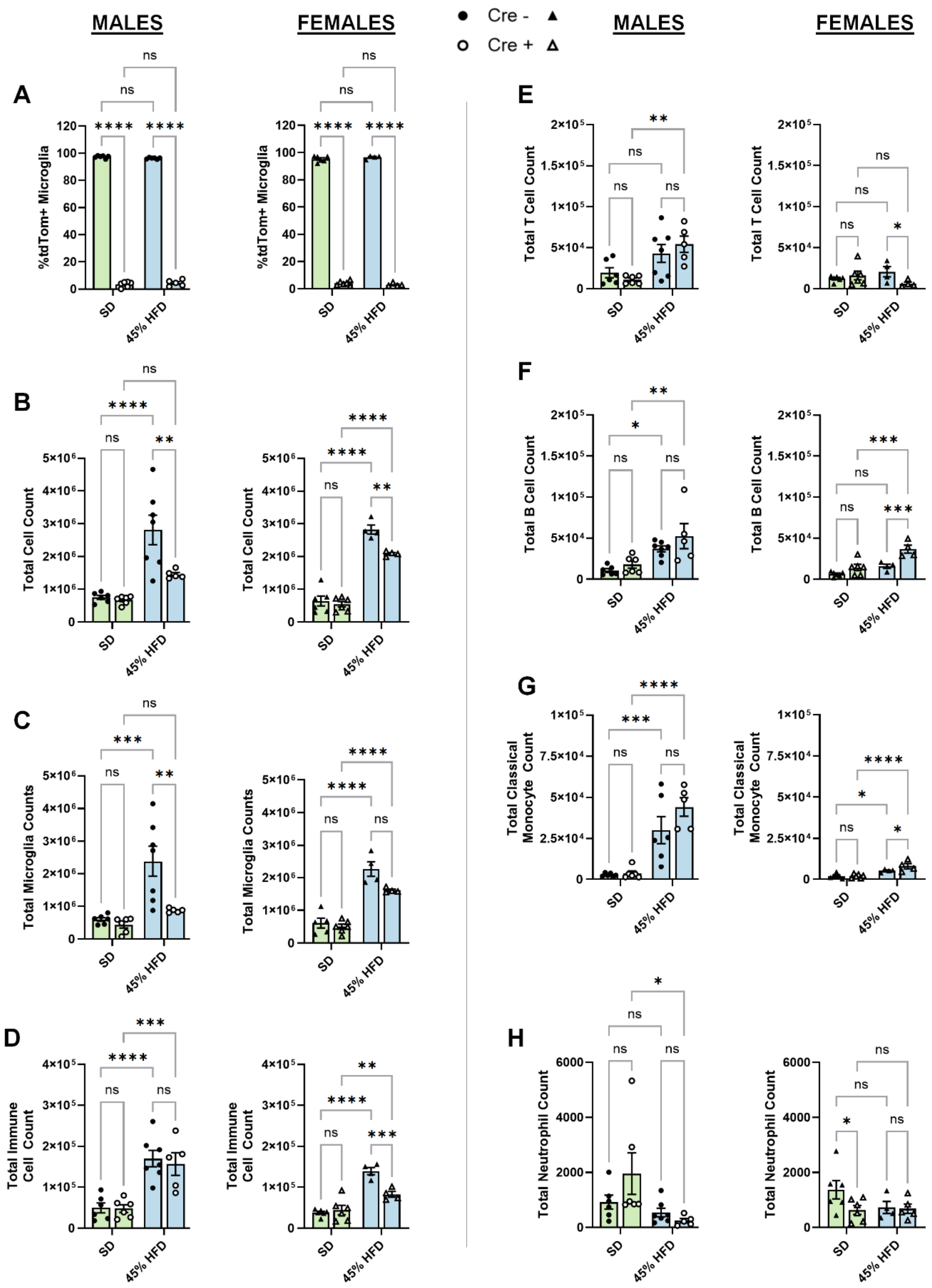


**Supplemental Figure 7. Flow cytometry characterization of immune cell compartment changes as a consequence of HFD feeding and microglial C3aR deletion. A)** Validation of successful retention/deletion of microglial tdTom-C3aR1 observed by percentage of tdTomato expressing microglia across male/female animals on SD or 45% HFD. **B)** Total cell count of immune cell preps ($CD45^{int-hi}$; microglia and all other immune cells) isolated from individual whole brains, showing elevated immune cell presence due to HFD that is rescued in absence of microglial C3aR. **C)** Total microglial counts, showing diet induced increase that is mitigated in absence of C3aR1, as is consistent with histology. **D)** Total immune cell ($CD45^{hi}$ cells; excluding $CD45^{Int}$ microglia), consisting of T cells (TCRβ+; **E**), B cells (CD19+; **F**), classical monocytes (Ly6C+ CD11b+; **G**), and neutrophils (Ly6G+ CD11b+; **H**). There was a diet-induced increase in overall between males and females, with subtle differences between sexes/genotypes in the various specific immune cells compartments. Error bars represent standard error of the mean (SEM).

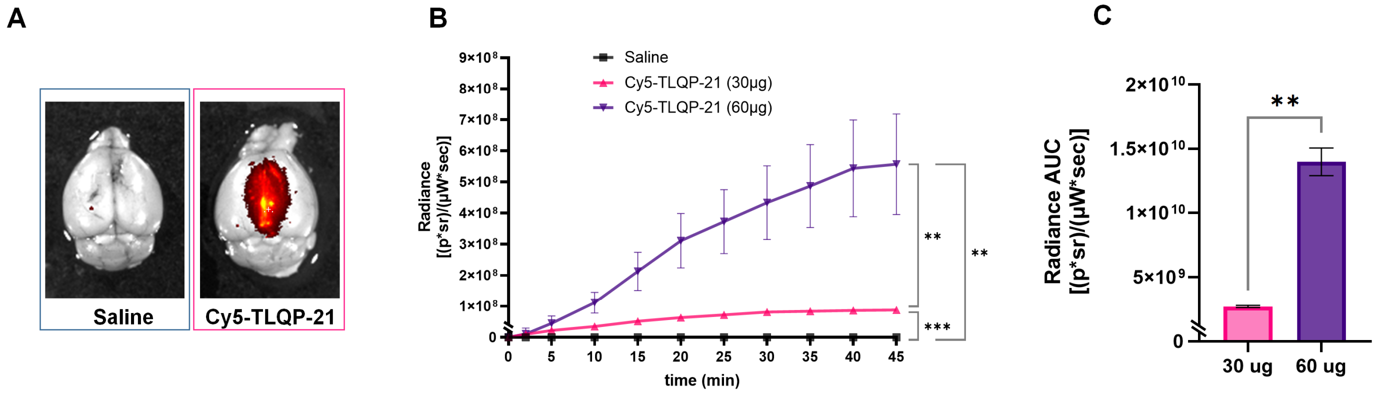


**Supplemental Figure 8. Cy5-conjugated TLQP-21 bypasses the blood-brain barrier via intranasal administration, as detected by in vivo imaging. A)** Whole brain ex-vivo imaging of mice administered saline (left panel) or Cy5-TLQP-21 (30 µg; right panel) 60 minutes using ROI capture. **B)** Time-course showing in-vivo uptake of Cy5-TLQP-21 over the first 45 minutes following administration of 30 µg (pink trace) or 60 µg (purple trace), with significant differences from saline-administered mice and between doses [one-way repeated-measures ANOVA ($F_{(1.002,10.02)}$ = 18.11, p = 0.0017, **p < 0.01, ***p < 0.001, n=3 mice per group)]. **C)** Area under the curve (AUC) comparison of 30 µg versus 60 µg time-courses. A significant difference in AUC between groups was quantified using unpaired t-test ($F_{(2,2)}$ = 177.8, p = 0.0087, **p < 0.01). ** indicates p<0.01, *** indicates p<0.001. Error bars represent standard error of the mean (SEM).

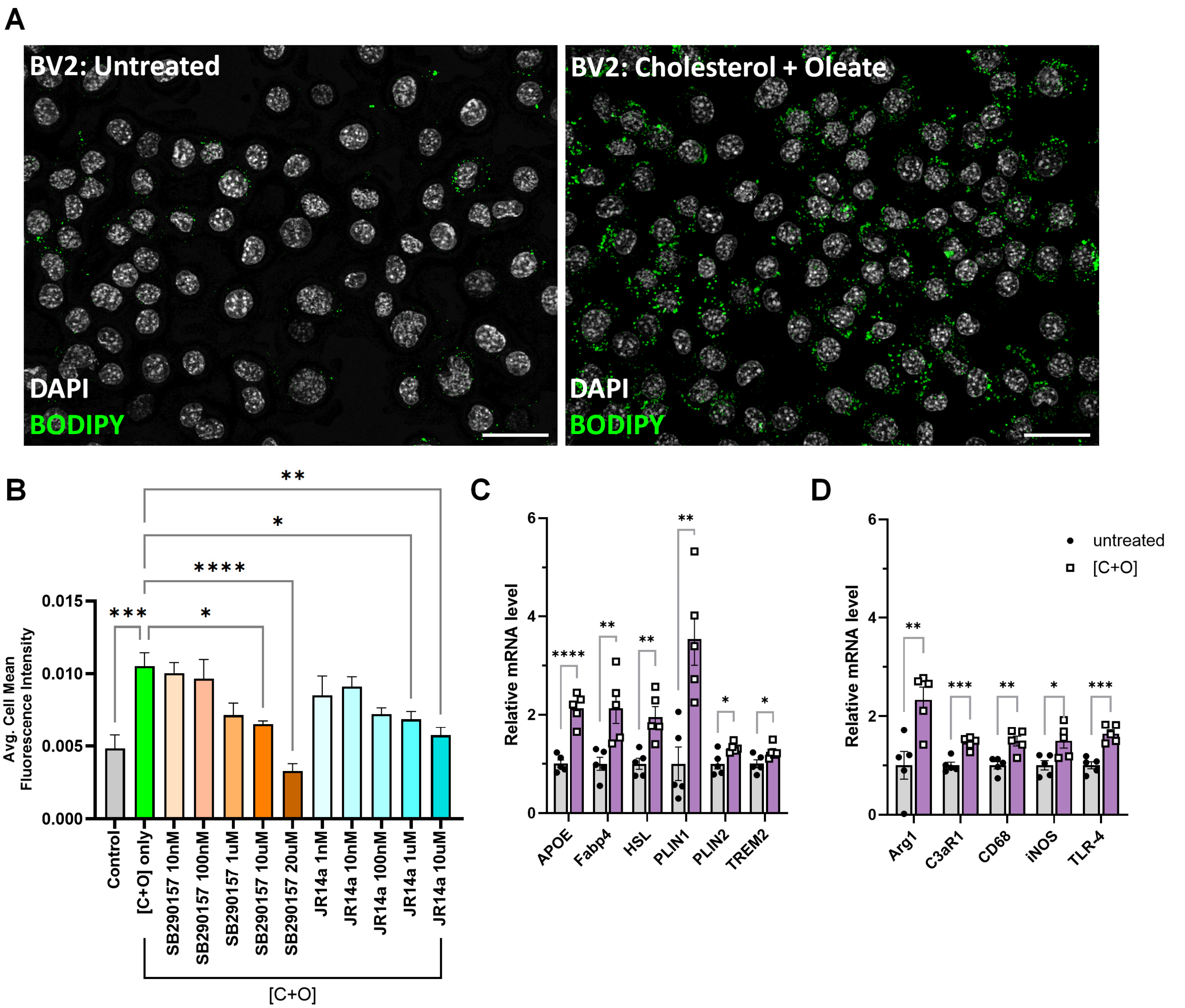


**Supplemental Figure 9. Effect of cholesterol + oleic acid on lipid loading and gene expression in BV2 microglia. A)** Representative 60x fluorescent images showing lipid droplets in DAPI/BODIPY stained BV2 cells either untreated or treated with Chol. + OA [C+O] for 18 hours. Scale bars are representative of 30μm. **B)** Quantification showing lipid loaded BV2 cells pre-treated with various concentrations of SB290157 or JR14a as a means of preventing [C+O]-induced lipid droplet accumulation. 1-way ANOVA followed by Tukey multiple comparisons test. * indicates p<0.05, ** indicates p<0.01, **** indicates p<0.0001. Relative expression of genes relevant for lipid metabolism (**D**) and inflammatory activation (**E**) in the context of lipid loading with C+O treatment normalized to untreated cells. Error bars represent standard error of the mean (SEM).

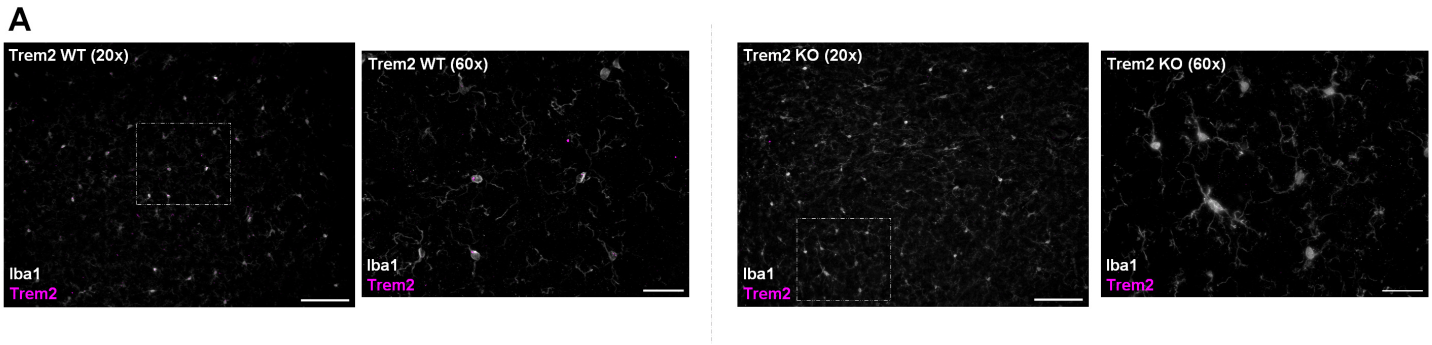


**Supplemental Figure 10. Validation of Trem2 immunofluorescent staining. A)** Representative 20x and 60x fluorescent images showing Trem2 signal present and colocalizing with hypothalamic microglia in brains collected from wildtype mice, but not that of germline, whole body Trem2 KO mice. Scale bars represent 100μm in the 20x image and 30μm in the 60x images.

**Supplementary Table 1: Change in Body Weights During 5-Day Tamoxifen Regimen**

| Diet | Sex | Genotype | Day 1 Avg. BW (g) | Day 5 Avg. BW (g) | n |
|---|---|---|---|---|---|
| *SD* | Male | C3aR1$^{lox/lox}$-CreER- | 32.47±1.65 | 32.33±1.59 | 12 |
| | Male | C3aR1$^{lox/lox}$-CreER+ | 31.01±1.33 | 30.91±1.33 | 10 |
| *SD* | Female | C3aR1$^{lox/lox}$-CreER- | 21.30±0.69 | 21.32±0.68 | 10 |
| | Female | C3aR1$^{lox/lox}$-CreER+ | 21.94±0.45 | 21.90±0.44 | 10 |
| *45% HFD* | Male | C3aR1$^{lox/lox}$-CreER- | 30.68±0.77 | 31.08±0.88 | 12 |
| | Male | C3aR1$^{lox/lox}$-CreER+ | 30.16±0.72 | 30.99±0.70 | 12 |
| *45% HFD* | Female | C3aR1$^{lox/lox}$-CreER- | 21.20±0.25 | 22.43±0.31 | 11 |
| | Female | C3aR1$^{lox/lox}$-CreER+ | 21.93±0.56 | 22.95±0.55 | 12 |
| *60% HFD* | Male | C3aR1$^{lox/lox}$-CreER- | 20.22±0.77 | 21.66±0.86 | 5 |
| | Male | C3aR1$^{lox/lox}$-CreER+ | 24.66±1.59 | 26.74±1.46 | 5 |
| *60% HFD* | Female | C3aR1$^{lox/lox}$-CreER- | 17.46±0.45 | 18.38±0.36 | 5 |
| | Female | C3aR1$^{lox/lox}$-CreER+ | 17.12±0.46 | 18.20±0.39 | 5 |

**Supplementary Table 2: Sequences of primers used in qPCRs**

| Gene | Forward | Reverse |
|---|---|---|
| *18S* | CGGGTGCTCTTAGCTGAGTGTCCCG | CTCGGGCCTGCTTTGAACAC |
| *Actin* | GGCACCACACCTTCTACAATG | GGGGTGTTGAAGGTCTCAAAC |
| *APOE* | GACCCTGGAGGCTAAGGACT | AGAGCCTTCATCTTCGCAAT |
| *Arg1* | AGCCAATGAAGAGCTGGCTGGT | AACTGCCAGACTGTGGTCTCCA |
| *C3aR1* | TCGATGCTGACACCAATTCAA | TCCCAATAGACAAGTGAGACCAA |
| *CD68* | TTCTGCTGTGGAAATGCAAG | AGAGGGGCTGGTAGGTTGAT |
| *Fabp4* | TGAAATCACCGCAGACGACAGG | GCTTGTCACCATCTCGTTTTCTC |
| *HSL* | GCTGGTGCAGAGAGACAC | GAAAGCAGCCGCCACGCG |
| *iNOS* | TGACGCTCGGAACTGTAGCAC | TGATGGCCGACCTGATGTT |
| *PLIN1* | AGAGGAGACAGATGAGGAGGAAG | AGATGGTGTTCTGCAGAGTCTTC |
| *PLIN2* | GACAGGATGGAGGAAAGACTGC | GGTAGTCGTCACCACATCCTTC |
| *TLR-4* | GAGGCAGCAGGTGGAATTGTAT | TTCGAGGCTTTTCCATCCAA |
| *TREM2* | GACCTCTCCACCAGTTTCTCC | TACATGACACCCTCAAGGACTG |

**Supplementary Table 3: Antibodies**

| Antibody | Source | Catalog number | Dilution |
|---|---|---|---|
| Guinea Pig anti NeuN | Millipore | ABN90 | 1:1000 |
| Rabbit anti Iba1 | Wako | 019-19741 | 1:1000 |
| Rabbit anti GFAP | Millipore | G9269 | 1:500 |
| Rat anti CD68 | Bio-Rad | MCA1957 | 1:300 |
| Sheep CSF R/CD115 | R&D Systems | AF3818 | 1:300 |
| Donkey anti Rabbit AlexaFluor 488 | Jackson Immuno | 711-545-152 | 1:500 |
| Donkey anti Rabbit AlexaFluor 647 | Jackson Immuno | 711-605-152 | 1:500 |
| Donkey anti Guinea Pig AlexaFluor 488 | Jackson Immuno | 706-545-148 | 1:500 |
| Donkey anti Rat AlexaFluor 647 | Jackson Immuno | 703-545-155 | 1:500 |
| Donkey anti Sheep Alexa Fluor 488 | Jackson Immuno | 713-545-003 | 1:500 |

**Supplementary Table 4: RNAscope probes**

| Probe | Source | Catalog number | Accession # |
|---|---|---|---|
| *Aldh1l1* | ACD Bio | 405891-C3 | NM_027406.1 |
| *C3aR1* | ACD Bio | 476751-C1 | NM_009779.2 |
| *Rbfox3* | ACD Bio | 313311-C3 | NM_001039167.1 |
| *tdTomato* | ACD Bio | 317041-C2 | LC311026.1 |
| *Tmem119* | ACD Bio | 472901-C3 | NM_146162.2 |

**Supplementary Table 5: Antibodies for Flow Cytometry (Brain & Small Intestine)**

| Antibody/Dye (Clone) | Source | Catalog number | Dilution |
|---|---|---|---|
| anti-CD45 BV785 (clone: 30-F11) | Biolegend | 103149 | 1:200 |
| anti-CD11b APC Fire 750 (clone: M1/70) | Biolegend | 101262 | 1:200 |
| anti-CX3CR1 BV605 (clone: SA011F11) | Biolegend | 149027 | 1:200 |
| anti-Ly6C PerCP-Cy5.5 (clone: HK1.4) | Biolegend | 128012 | 1:200 |
| anti-Ly6G Pac Blue (clone: 1A8) | Biolegend | 127612 | 1:200 |
| anti-CD206 PE-Cy7 (clone: C068C2) | Biolegend | 141720 | 1:200 |
| anti-TCRβ APC (clone: H57-597) | Biolegend | 109212 | 1:200 |
| anti-CD19 FITC (clone: 6D5) | Biolegend | 115506 | 1:200 |
| Ghost Dye Violet 510 Viability Dye | Tonbo Biosciences | 13-0870-T100 | 1:1000 |

| Antibody/Dye (Clone) | Source | Catalog number | Dilution |
|---|---|---|---|
| anti-Timd4 APC (clone: RMT4-54) | Biolegend | 13008 | 1:100 |
| anti-CD192 (Ccr2) BV421 (clone: SA203G11) | BD Biosciences | 150605 | 1:100 |
| anti-CD206 PE/Cy7 (clone: C068C2) | Biolegend | 141720 | 1:100 |
| anti-MHCII PerCP-CY5.5 (clone: M5/114.15.2) | Biolegend | 127612 | 1:100 |
| anti-CD64 BV421 (clone: X54-5-7.1) | Biolegend | 139309 | 1:100 |
| anti-CD11b BV510 (clone: M1/70) | Biolegend | 101245 | 1:100 |
| anti-F4/80 BV650 (clone: BM8) | Biolegend | 123149 | 1:100 |
